# TALEN-mediated intron editing of HSPCs enables transgene expression restricted to the myeloid lineage

**DOI:** 10.1101/2024.03.05.583596

**Authors:** Eduardo Seclen, Jessica C. Jang, Aminah O. Lawal, Sylvain Pulicani, Alex Boyne, Diane Tkach, Alexandre Juillerat, Philippe Duchateau, Julien Valton

**Author notes:** Correspondence should be addressed to J.V.

## Abstract

Gene therapy in hematopoietic stem and progenitor cells (HSPCs) shows great potential for the treatment of inborn metabolic diseases. Typical HSPC gene therapy approaches rely on constitutive promoters to express a therapeutic transgene, which is associated with multiple disadvantages. Here, we propose a novel promoter-less intronic gene editing approach that triggers transgene expression only after cellular differentiation into the myeloid lineage. We integrated a splicing-competent eGFP cassette into the first intron of *CD11b* and observed expression of eGFP in the myeloid lineage but minimal to no expression in HSPCs or differentiated non-myeloid lineages. *In vivo*, edited HSPCs successfully engrafted in immunodeficient mice and displayed transgene expression in the myeloid compartment of multiple tissues. Using the same approach, we expressed alpha-L-iduronidase (IDUA), the defective enzyme in Mucopolysaccharidosis type I, and observed a 10-fold supraendogenous IDUA expression exclusively after myeloid differentiation. Edited cells efficiently populated bone marrow, blood, and spleen of immunodeficient mice, and retained the capacity to secrete IDUA *ex vivo*. Importantly, cells edited with the eGFP and IDUA transgenes were also found in the brain. This approach may unlock new therapeutic strategies for inborn metabolic and neurological diseases that require the delivery of therapeutics in brain.

## Introduction

Gene editing in hematopoietic stem and progenitor cells (HSPCs) has enabled the treatment of multiple previously uncurable genetic diseases. Edited therapeutic HSPCs can engraft in the patient’s bone marrow, self-replicate, differentiate and populate other hematopoietic organs, propagating the therapeutic effects systemically and indefinitely after a single intervention. This is the basis of the HSPC gene therapy for many hematological and metabolic genetic diseases^1^. In addition, a subset of HSPC can differentiate into cells able to cross the blood brain barrier and become microglia, propagating the therapeutic effects into the brain. Neurodegeneration is a common trait of many genetic metabolic diseases, which makes HSPC gene therapy a better therapeutic alternative than enzymatic replacement therapy. Beyond metabolic diseases, edited HSPC could act as Trojan horses for the delivery of genetically-encoded therapeutics in the brain to address neurodegenerative conditions like Parkinson’s^2^.

Common therapeutic modifications in HSPC include the correction of single point mutations, gene inactivation or, more frequently, the integration of an expression cassette to provide the patient with a therapeutic protein *in trans*. Most of the clinically advanced therapeutic HSPC approaches addressing inherited metabolic diseases are based on the integration of an expression cassette into a random region of the genome using lentiviral vectors^3,4,5^. Editing HSPC with lentiviral vectors have shown to be efficient and safe, however, while the latest generations of lentiviral vector are engineered to reduce the risk of insertional mutagenesis, this type of event still occur, as observed in recent clinical trials^6^. The development targeted nucleases like TALEN or CRISPR Cas9 has reduced the risk of insertional mutagenesis given its ability to assist the integration of an expression cassette into a specific genome site, commonly known as safe harbor locations^7^. Using a targeted or safe harbor approach a second generation of treatments for metabolic diseases could be on its way given the positive results obtained in many preclinical studies^8^.

However, both lentiviral and targeted integration approaches generally rely on the use of ubiquitous promoters to drive transgene expression. Two main disadvantages are derived from the use of ubiquitous promoters: the potential dysregulation of nearby loci^9^ and the potential detrimental effect of transgene’s expression at the stem cell level for some transgenes^10,11^. In addition, ubiquitous promoters lead to expression of the transgene in all HSPC-derived differentiated lineages. While this strategy could be beneficial to address diseases that result from the systemic absence of a given protein or enzyme, it could be unnecessary or even, detrimental, to address other diseases that result from a protein deficiency in a single lineage. This is the case of beta-globin in sickle cell disease^12^ or certain immunodeficiencies mostly affecting lymphoid cells like RAG1 deficiency^13^. In addition, erythroid, myeloid, and lymphoid lineages distribute differently in multiple organs after engraftment. For example, B cells are highly prevalent in the bone marrow, T cells in lymph nodes, while myeloid and erythroid lineages have a more systemic distribution. Enabling transgene expression exclusively in certain lineages or cell types could more efficiently address certain genetic conditions.

In this paper, we developed a promoterless, intron-specific gene insertion approach for HSPC, that restricts the expression of a desired transgene to a specific lineage, preventing overexpression at the stem cell level or in other differentiated lineages. We focused on the myeloid lineage given its systemic distribution and potential to reach the brain as HSPC-derived microglia, which could allow enable therapies for both metabolic and neurological disorders. Specifically, we inserted a splicing competent promoter-less cassette into the first intron of the *CD11b* myeloid gene and allowed the RNA splicing machinery to direct the expression of the inserted transgene without affecting endogenous *CD11b*. Using the enhanced green fluorescent protein (eGFP) as a reporter transgene, we found that transgene expression was restricted to the myeloid compartment after *in vitro* differentiation and after *in vivo* engraftment into two immunodeficient mouse models. We further demonstrate the translatability of our gene insertion approach by inserting alpha-L-iduronidase (IDUA), the deficient enzyme in Mucopolysaccharidosis type I (MPS-I). In addition to displaying myeloid lineage specific expression of IDUA *in vitro*, we found that edited HSPC exhibited robust engraftment in the bone marrow, displayed multi-lineage differentiation in various hematopoietic tissues and showed significant presence in the brain. Furthermore, edited cells retrieved from bone marrow retained their capacity to secrete IDUA *ex vivo*. Given its unique characteristics, we believe this HSPC editing technology could be disruptive in the development of therapeutics for many metabolic and neurological diseases.

## Results

### Editing HSPCs at the *CD11b* intron enables specific transgene expression in the myeloid compartment

Our goal was to edit HSPCs and induce the expression of a transgene exclusively in the myeloid lineage. Our editing strategy relied on the integration of a transgene into the first intron of *CD11b* (Fig 1a). As a first proof of concept, we designed a transgene encoding eGFP linked to the first *CD11b* exon by a 2A self-cleavable peptide. This expression cassette was designed with homology arms to drive gene insertion via homologous recombination and was flanked by splice acceptor and donor sequences to allow the transcription of a polycistronic eGFP-CD11b mRNA molecule. This design was chosen so that transgene expression was linked to that of *CD11b* and triggered only after myeloid differentiation of edited HSPC. As a control for ubiquitous transgene expression in HSPCs and all differentiated hematopoietic lineages, we inserted an eGFP expression cassette carrying the phosphoglycerate kinase (PGK) promoter into the AAVS1 safe harbor locus, referred as Ctrl-PGK hereafter.

**Figure 1.**
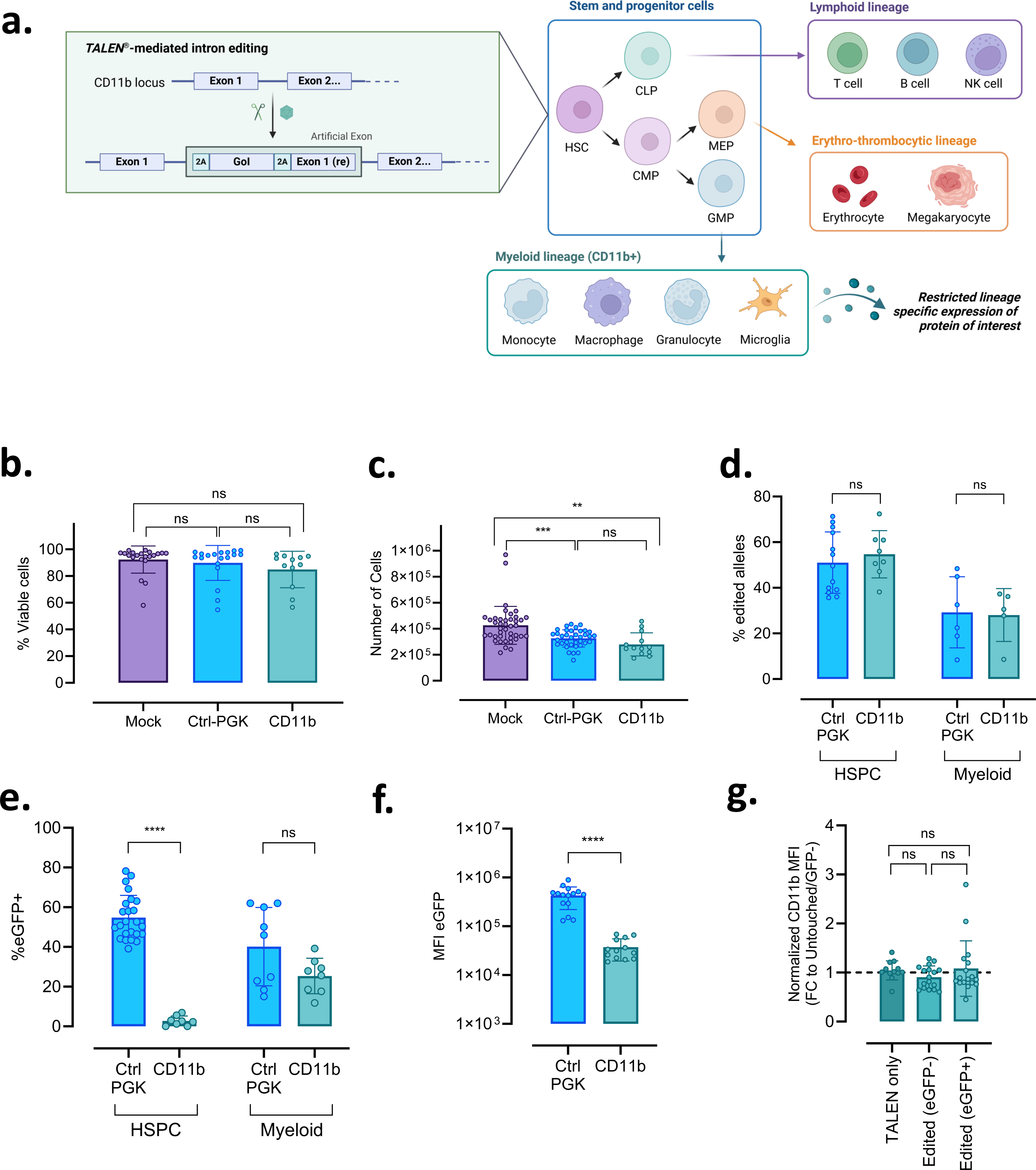
**Editing HSPCs at the *CD11b* intron enables specific transgene expression in the myeloid compartment**. **a.** Diagram depicting the TALEN®-mediated intron gene editing strategies and subsequent cellular expression upon differentiation. GoI; gene of interest; HSC; hematopoietic stem cell, CLP; common lymphoid progenitor, CMP; common myeloid progenitor, MEP; megakaryocyte-erythroid progenitor, GMP; granulocyte-macrophage progenitor. **b.** Viability measured by the proportion of 7AAD-AnnexinV- cells measured by flow cytometry 24 hours after gene editing. **c.** Number of viable cells 24 hours after gene editing. **d.** Allelic editing rates measured 5 days after editing in HSPC media or 14 days after editing and myeloid differentiation by digital PCR. **e.** Editing rates measured by flow cytometry as proportion of eGFP+ cells 5 days after editing in HSPC media (ungated) or 14 days after editing and myeloid differentiation (gated in CD14high cells). **f.** Intensity of expression of eGFP from each editing strategy measured by mean fluorescence intensity (MFI) of eGFP+ cells, gated in CD14high cells 14 days after gene editing and myeloid differentiation. **g.** Mean fluorescence intensity of *CD11b* expression measured 14 days after gene editing and myeloid differentiation in TALEN only treated samples (eGFP-), CD14high/eGFP- cells (unedited or with intronic indels), and CD14high/eGFP+ cells (edited), normalized to values from unedited differentiated cells. Data are represented as mean ± standard deviation. Statistical comparisons were performed using unpaired T tests when comparing two groups (1d, 1e, 1f), or one way ANOVA when comparing three groups followed by a Tukey’s Honestly-Significant Difference post-hoc test between each two groups (1b, 1c, 1g); ns= non-significant, **p≤0.01, ***p≤0.001, ****p≤0.0001.

Our HSPC gene editing protocol, relying on TALEN mRNA electroporation followed by adeno-associated virus transduction, resulted in viability in the 85-90% range (Fig 1b), with 66- 76% of mock electroporated cells preserved at 24h after editing (Fig 1c). Allelic editing rates for *CD11b* were 55% (Fig 1d), in similar range to the Ctrl-PGK control in undifferentiated HSPC.

Our *CD11b* intron editing-based approach was designed to insert a transgene into HSPCs and delay its expression until after myeloid differentiation. To investigate that aspect, we subjected a fraction of the edited cells to myeloid differentiation for up to 21 days. After myeloid differentiation *in vitro*, HSPCs lost the CD34 marker (with 98% CD34+ before differentiation vs. 4% after differentiation) and gained the expression of multiple myeloid markers including CD14 (63%), CD15 (65%), CD11b (88%), S100A9 (79%), and CD68 (70%, Fig S1). By design, in Ctrl-PGK edited cells, we observed eGFP expression before (average of 55% eGFP+ cells, Fig 1e), and after (40% eGFP+) myeloid differentiation. In stark contrast, HSPCs edited at the *CD11b* locus barely expressed any eGFP in the undifferentiated state (<3%), despite similar levels of allelic gene editing when compared with the Ctrl-PGK condition (Fig 1d and 1e). In HSPCs edited at the *CD11b* locus, eGFP expression was only induced after myeloid differentiation, with 25% of cells expressing eGFP within the myeloid compartment (Fig 1e). Mean fluorescence intensity values of eGFP+ were higher when its expression was driven by the PGK promoter in comparison to the *CD11b* endogenous promoter, as expected (4.3e5 vs 3.7e4 units, respectively, Fig 1f). These results indicate that TALEN-mediated *CD11b* intron editing in HSPCs restricts transgene expression to the myeloid lineage.

Our transgene insertion approach was designed to be non-disruptive, aiming to avoid inducing any changes in the expression of the targeted *CD11b* gene. To confirm this, we compared *CD11b* expression levels in differentiated HSPCs after several treatment conditions (unedited or *CD11b* TALEN-treated, in the absence or presence of AAV donor template). We observed similar levels of *CD11b* expression in cells treated with TALEN without AAV (i.e., harboring indels in the *CD11b* intron) and cells treated with TALEN and AAV (Fig 1g). More specifically, the eGFP+ (successfully edited) and eGFP- (mixture of unedited and indel-containing cells) fractions of the edited cells displayed the same mean fluorescence intensity for *CD11b*. These data suggest that our editing approach does not affect the endogenous expression of the edited locus.

### HSPCs edited at the *CD11b* intron maintain their differentiation potential *in vitro* and secrete IDUA enzyme after myeloid differentiation

After demonstrating that the intronic gene editing approach is feasible, we assessed its potential to overexpress a protein of interest. To do so, we focused on MPS-I, a lysosomal storage disease (LSD) resulting from deficiencies in the IDUA enzyme. We began by optimizing editing rates for inserting an IDUA cassette into *CD11b* with a similar editing to the one described above (Fig 2a). Edited HSPCs were subject to bulk myeloid differentiation to assess the secretion of IDUA *in vitro*. Edited cells were also subjected to clonal myeloid differentiation using a colony forming unit (CFU) assay. This allowed us to quantify the myeloid differentiation potential of edited HSPC as well as to determine clonal editing rates of our editing approach (Fig 2a). Consistent with the data obtained for the eGFP transgene, the IDUA transgene had a minimal (but statistically significant) effect on HSPC viability and remaining cells (Fig 2b and 2c). The frequencies of IDUA transgene insertion into the *CD11b* locus of undifferentiated HSPCs averaged 56% (Fig 2d).

**Figure 2.**
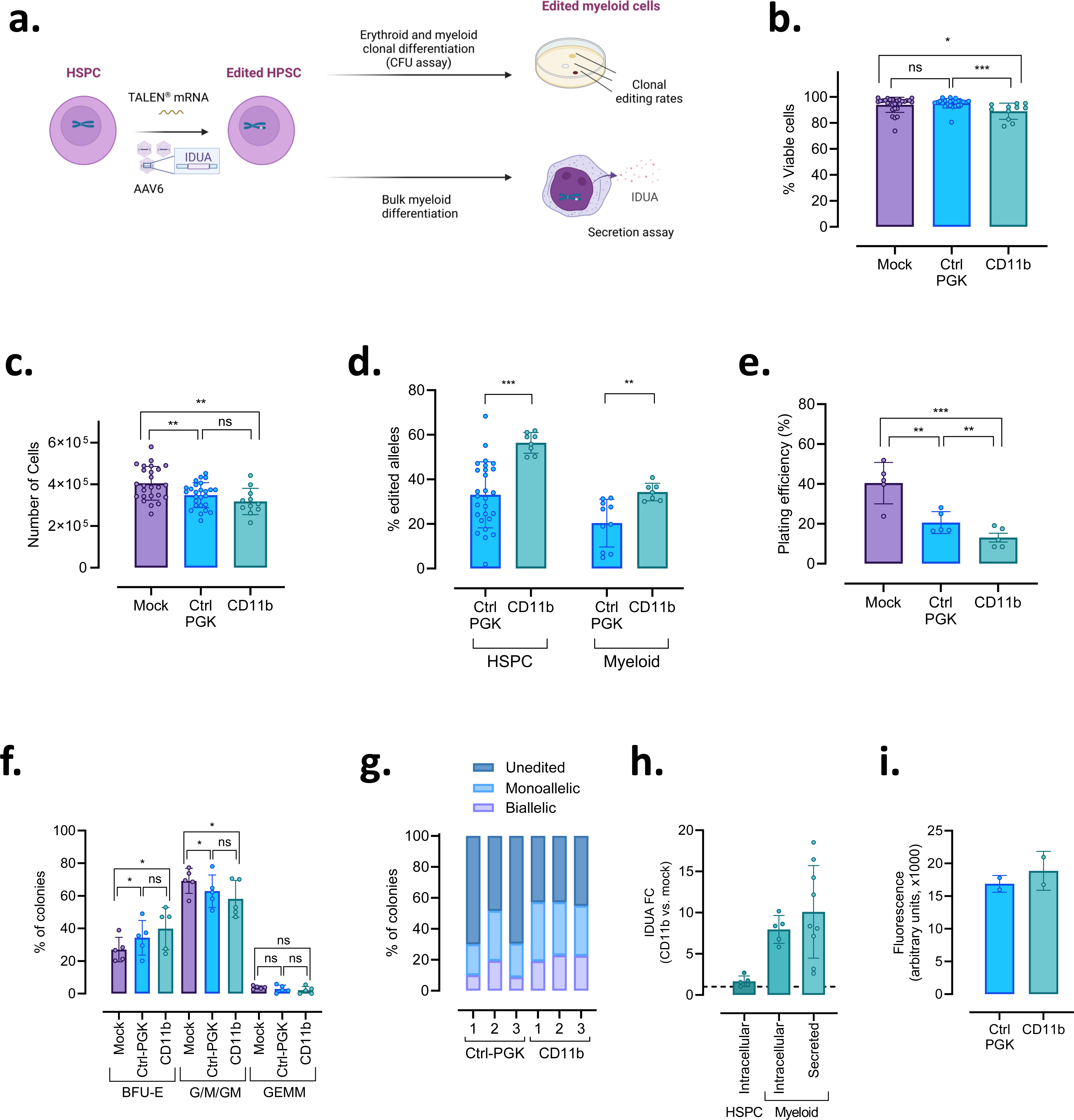
HSPCs edited at the *CD11b* intron maintain their differentiation potential *in vitro* and secrete IDUA enzyme after myeloid differentiation**. a.** Diagram depicting editing process and assay readout after cell differentiation. **b.** Viability measured as the proportion of 7AAD- Annexin V- cells measured by flow cytometry 24 hours after gene editing. **c.** Number of viable cells 24 hours after gene editing. **d.** Editing rates measured 5 days after editing in HSPC media or 14 days after editing and myeloid differentiation by digital PCR. **e.** Plating efficiency of CFU assay, measured as the percentage of seeded cells that successfully generated a colony in the methylcellulose plate. **f.** Percentage of the distinct types of colonies in CFU assay. Erythroid burst forming units (BFU-E), Granulocyte/Macrophage/Granulocyte-Macrophage (G/M/GM), or the more primitive granulocyte-erythroid-monocyte-megakaryocyte (GEMM). **g.** Editing rates in clones isolated from CFU plates (n=3 donors). **h.** Normalized IDUA levels determined by ELISA in cell lysates of undifferentiated and cell lysates and supernatants for differentiated *CD11b*-IDUA edited HSPC. **i.** Enzymatic activity of 15 ng of IDUA obtained from the supernatants of Ctrl-PGK (unmodified IDUA enzyme) or *CD11b* edited cells (IDUA with extra amino- and carboxi- residues, product of the *CD11b* intron editing strategy). Data are represented as mean ± standard deviation. Statistical comparisons were performed using unpaired T tests when comparing two groups (fig 2d), or one way ANOVA when comparing three groups followed by a Tukey’s Honestly-Significant Difference post-hoc test between each two groups (fig 2b, 2c, 2e, 2f); ns= non-significant, *p≤0.05, **p≤0.001, ***p≤0.001.

As with the eGFP expression approach above, this editing method relies on the mRNA splicing machinery to correctly express the transgene and the endogenous gene targeted. To characterize the integration of the transgene at the genomic level and the splicing process at the mRNA level, we performed long read clonal sequencing on genomic DNA and fully spliced mRNA. Our results confirmed that the entire IDUA transgene was integrated between the first and second exons of *CD11b* (Fig S2). To characterize the final chimeric messenger RNA molecule, we isolated messenger RNA molecules from edited HSPCs subjected to myeloid differentiation, generated the *CD11b* specific complementary DNA (cDNA), amplified the region comprising the first and second exons of *CD11b*, and characterized the transcripts using long-read sequencing. In unedited cells, we found the expected sequence including the first two exons of *CD11b* (Fig S3). In edited cells, in addition to the unedited *CD11b* sequences, we found the intended bi-cistronic IDUA-*CD11b* mRNA molecule, suggesting that our insert was successfully processed by the RNA machinery and integrated into the *CD11b* mRNA. Importantly, no additional intronic sequences were observed upstream or downstream of the inserted cassette, suggesting that the splicing process was precise.

In addition to efficient gene editing and successful splicing of the IDUA transgene, our goal was to maintain differentiation capacity of edited HSPCs and elicit a broader, potentially systemic effect. We therefore assessed the capacity of edited HSPCs to differentiate *in vitro* using a CFU assay. We observed that mock-treated HSPCs had greater plating efficiency, defined as the proportion of plated cells that develop into a cellular colony (averaging 40%) than did the Ctrl-PGK (21%) or *CD11b* edited (13%) HSPCs (Fig 2e). Compared to unedited cells, edited HSPCs developed a higher proportion of erythroid progenitors (mean of 27% vs. 34%/40%, respectively, Fig 2f), and a lower proportion of granulocyte/monocyte progenitors (mean values of 69% vs. 63%/58%). While these differences were statistically significant, the effect sizes were small. In addition, no differences were observed in the erythroid or granulocyte/monocyte populations in HSPCs edited at *CD11b* locus compared to Ctrl-PGK cells. This suggests that editing at the *CD11b* intron did not negatively impact the myeloid differentiation process.

Editing efficiency is usually reported like bulk allelic editing rates. However, this metric underestimates the actual proportion of edited cells (edited with the IDUA transgene at one or two alleles) that can elicit a therapeutic benefit. CFU assays allowed us to characterize the proportion of mono- and bi-allelically modified cells via ddPCR analysis of individual colonies. For HSPCs edited at the *CD11b* locus, the average bulk allelic editing rates of differentiated cells in bulk liquid culture from multiple experiments was 35% (Fig 2d), while the proportion of successfully edited colonies in the semi solid CFU assay was 56% (Fig 2g). Colonies that were edited at one allele were more frequent than those edited at both, as expected (62% vs. 38%, respectively).

Finally, we evaluated the ability of our intron edited HSPC to generate supra-endogenous levels of IDUA. After editing and differentiating HSPCs *in vitro* into the myeloid lineage, we quantified the amount of intracellular and secreted IDUA by ELISA (Fig 2h). While IDUA presence in *CD11b* edited HSPC before differentiation was low, the expression (intracellular) and secretion (extracellular) of IDUA increased as much as 10- and 19-fold, respectively (average of 8- and 10-fold across multiple donors, respectively) in comparison with the endogenous IDUA secretion of unmodified healthy cells after myeloid differentiation, demonstrating that IDUA was expressed and secreted in a lineage-specific manner. In addition, we confirmed that the recombinant IDUA generated by our intron editing approach was as functional as one generated from Ctrl-PGK edited cells (Fig 2i) and that the extra amino acid residues at either the C- or N-terminal end of the IDUA derived from the use of 2A peptides (Fig 1a) did not negatively impact its enzymatic activity.

### HSPCs edited at the *CD11b* intron maintain their engraftment and differentiation capacity *in vivo*

We next evaluated the translatability of our intron-based gene editing approach *in vivo* by engrafting HSPCs edited at the *CD11b* locus with the eGFP or IDUA transgenes into 2 different mouse models: immunodeficient NSG (NOD.Cg-Prkdc^SCID^Il2rg^tm1Wjl^/SzJ) mice and SGM3 mice, a NSG strain further genetically engineered to express human cytokines (IL3, GM-CSF, and SCF) that better support myeloid differentiation^14,15^ (fig 3a). As a positive control for expression in all hematopoietic lineages we used again the insertion of a PGK expression cassette into the AAVS1 locus (Ctrl-PGK). In these mouse models, human HSPCs engraft into the bone marrow of the animal and generate both myeloid and lymphoid progeny that are able to populate other hematopoietic tissues.

**Figure 3.**
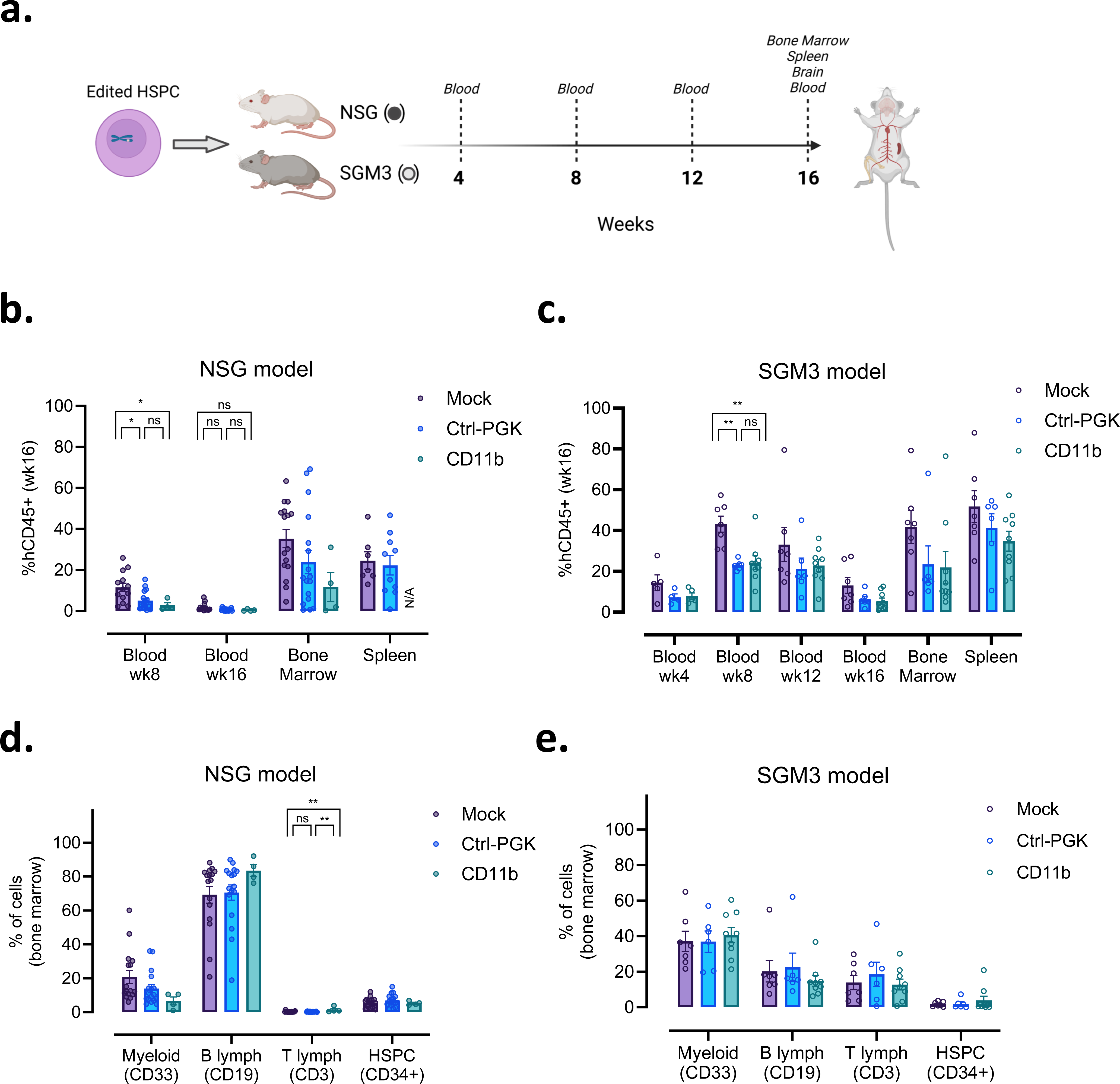
HSPCs edited at the *CD11b* intron maintain their engraftment and differentiation capacity *in vivo*. a. Diagram depicting timelines for *in vivo* experiments. **b.** Engraftment in blood, bone marrow and spleen in NSG cohorts, measured as the proportion of human cells (hCD45+). **c.** Engraftment in blood, bone marrow and spleen in SGM3 cohorts, measured as the proportion of human cells (hCD45+). **d.** Percentage of myeloid, B cell lymphoid, T cell lymphoid and HSPC in the bone marrow of NSG cohorts. **e.** Percentage of myeloid, B cell lymphoid, T cell lymphoid and HSPC in the bone marrow of SGM3 cohorts. N/A: Not analyzed. Empty circles = SGM3. Filled circles = NSG. Data are represented as mean ± standard deviation. Statistical comparisons were performed using one way ANOVA followed by a Tukey’s Honestly-Significant Difference post-hoc test between each two groups; Tukey’s p values are only shown if ANOVA was significant; ns= non-significant, *p≤0.05, **p≤0.01.

In the NSG mouse model, the lowest human chimerism was observed in blood, with averages of 7.3% and 1.1% at 8 and 16 weeks after injection, respectively (fig 3b). Level of chimerism in bone marrow and spleen were higher, ranging 10-35%. Overall, lower levels of human chimerism were observed in most tissues of animals injected with edited compared to those injected with unedited HSPCs, although differences were only statistically significant in blood. Importantly, there were no differences between the *CD11b* and Ctrl-PGK group in any tissue. In the SGM3 model, human chimerism was higher, ranging 20-50% in most hematopoietic tissues (fig 3c). Although not significant, a similar trend towards lower engraftment of edited cells was observed in most tissues. This reduced engraftment capability of edited HSPCs is common and has been reported previously^16–18^.

We then evaluated the differentiation capacity of HSPC and assessed whether our editing approach could affect this process in the bone marrow and other hematopoietic tissues including the blood and spleen. To do so, we characterized the presence of myeloid (CD33), B lymphoid (CD19), T lymphoid (CD3), and stem cell (CD34) lineages in the blood, spleen, and bone marrow (Fig 3d and 3e). In the bone marrow, a predominant B cell population was observed in NSG mice (fig 3d), while myeloid cells were more prevalent in the SGM3 model (fig 3e), likely due to the presence of human cytokines favoring myeloid differentiation in this model. In general, no statistical differences in the proportion of different lineages were observed among treatment groups in either the NSG or the SGM3 model. Of note, in the NSG model a trend towards a smaller proportion of myeloid cells and higher proportion of B cells was observed in the *CD11b* edited group compared with the unedited group. However, this was not statistically significant and not observed in the SGM3 model. The only statistically significant difference in bone marrow was a higher proportion of T cells in the SGM3 model for the *CD11b* condition, but numerical differences were negligible compared to the unedited or Ctrl-PGK condition (average < 1.5%). No differences were observed in lineage distribution among treatment groups in the spleen or blood (Fig S4). Altogether, these data indicated that our gene editing approach preserved the engraftment and differentiation potential of HSPCs.

### HSPCs edited at the *CD11b* intron persist *in vivo* and support myeloid-specific transgene expression

After confirming that editing protocols did not impact the engraftment and differentiation capacity of HSPCs, we evaluated the persistence of the editing signature in cells engrafted in the bone marrow and other hematopoietic tissues. Input *in vitro* editing frequencies of the eGFP-edited HSPCs batches tested *in vivo* are shown in Fig 4a ranging from 46% to 78%. Sixteen weeks after injection, animals were euthanized and the proportion of eGFP+ cells was analyzed in each lineage. In concordance with *in vitro* results, HSPC cells edited at the *CD11b* locus with an eGFP cassette injected into the SGM3 mouse model developed a myeloid-specific pattern of eGFP expression with eGFP levels of 5.4%, 8.2%, and 13.9% in the myeloid compartment of bone marrow, blood and spleen, respectively, with little to no expression in the HSPC, B cell, or T cell subfractions (fig 4b). This was in stark contract with the ubiquitous pattern of expression of SGM3 (fig 4c) or NSG (fig 4d) animals injected with HSPC edited with the PGK-eGFP cassette, which displayed a similar level of eGFP expression in all lineages analyzed in bone marrow, blood and spleen. Of note, 3.5% of cells from the T cell lymphoid lineage found in the bone marrow of SGM3 animals injected with *CD11b* edited HSPC were eGFP+. We then analyzed a subgroup of these samples for expression of *CD11b* (Fig S5) and found that only cells expressing *CD11b* were eGFP+. This data confirmed that our intron editing approach enabled to hijack the regulatory elements of *CD11b* to drive the expression of eGFP, which was generally restricted to the myeloid lineage *in vivo*.

**Figure 4.**
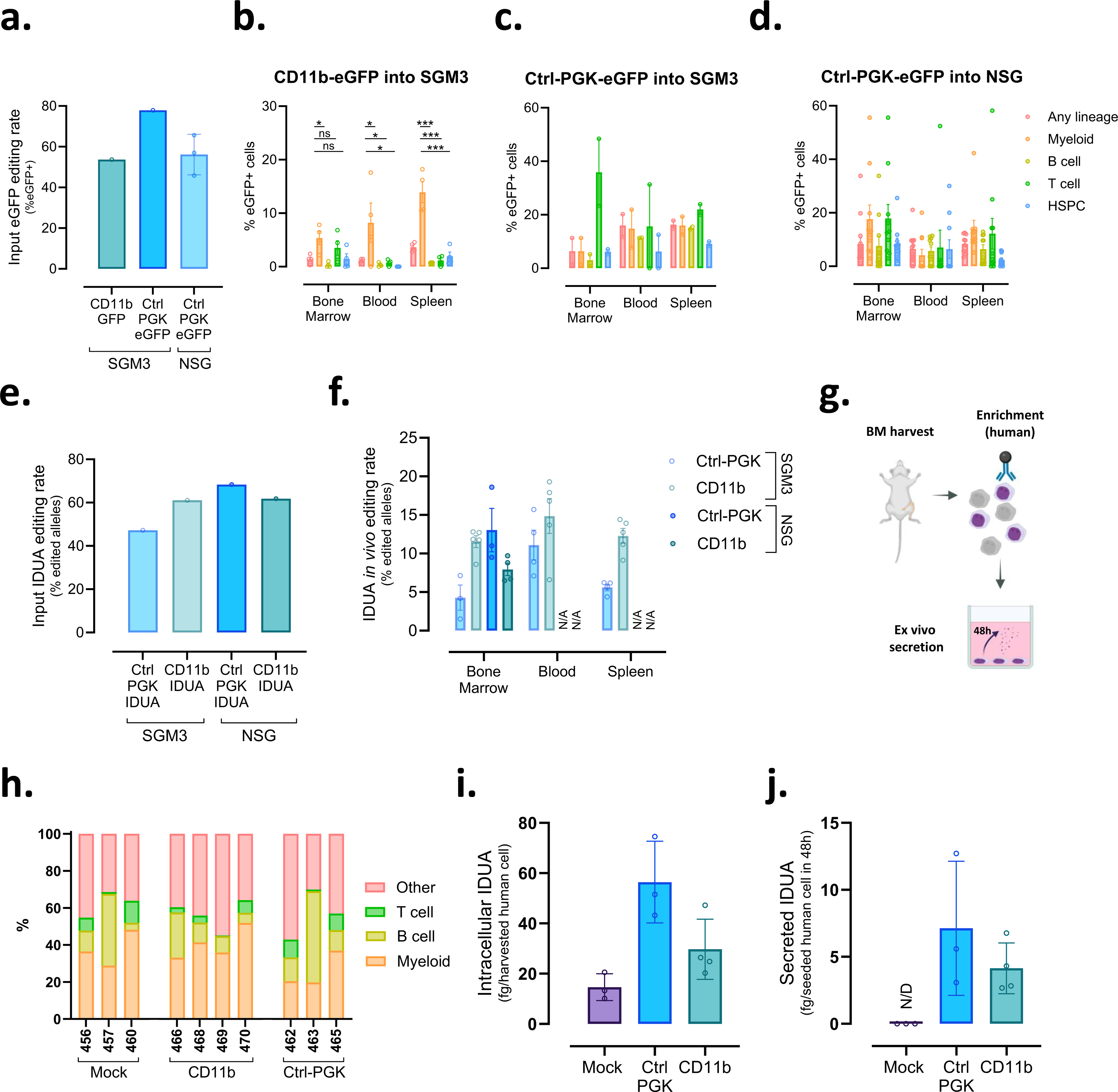
**HSPCs edited at the *CD11b* intron support myeloid-specific transgene expression *in vivo.* a**. Input editing rates of HSPC edited with eGFP cassettes and injected in NSG or SGM3 animals, as determined by eGFP expression measured 5 days after editing for the Ctrl-PGK group (ungated) or 14 days after myeloid differentiation for the *CD11b* group (gated in CD14high cells). **b.** Percentage of edited cells (eGFP+) within different lineages in the blood, bone marrow and spleen of SGM3 animals engrafted with *CD11b*-eGFP edited HSPCs, SGM3 animals engrafted with Ctrl-PGK-eGFP edited HSPCs (**c**), or NSG animals engrafted with Ctrl-PGK-eGFP edited HSPCs (**d**). **e**. Input allelic editing rates of HSPC edited with the IDUA cassette and injected in NSG or SGM3 animals, as determined 5 days post editing by ddPCR. **f.** Percentage of edited human alleles from cells recovered from blood, bone marrow and spleen of NSG and SGM3 animals engrafted with HSPCs edited with the IDUA cassette. **g.** Diagram depicting *ex vivo* IDUA secretion assay from bone marrow cells isolated from SGM3 animals injected with HSPC edited with the IDUA cassette. **h.** Lineage distribution in bone marrow of animals whose cells were used to set up the *ex vivo* IDUA secretion assay. **i.** Levels of IDUA detected by ELISA in the cell pellets of *ex vivo* cell cultures derived from the bone marrow of SGM3 animals **j.** Levels of IDUA detected by ELISA in the supernatants of *ex vivo* cell cultures derived from the bone marrow of SGM3 animals. N/A: Not analyzed. N/D: not detected. Empty circles = SGM3. Filled circles = NSG. Data are represented as mean ± standard deviation. Statistical comparisons were performed using one way ANOVA followed by a Tukey’s Honestly-Significant Difference post-hoc test between each two groups (fig4b, 4c, 4d). Tukey’s p values are only shown if ANOVA was significant; ns= non- significant, *p≤0.05, ***p≤0.001.

We also evaluated the engraftment of HSPCs edited with the IDUA transgene and injected into SGM3 and NSG animals. In these cohorts, the input *in vitro* allelic editing rates of injected HSPC ranged from 47% to 68% (fig 4e). *In vivo*, we found edited cells in the bone marrow of all NSG and SGM3 animals, with allelic editing rates ranging from 2% to 19% (Fig 4f). *In vivo* editing rates of SGM3 animals injected with Ctrl-PGK edited HSPC were lower than the rest in the bone marrow averaging 4.3%, with higher values in blood (11.1%) and spleen (5.6%); perhaps due to the lower input *in vitro* editing rate of HSPCs. These data demonstrated that HSPCs edited with the IDUA transgene can efficiently engraft in an immunodeficient mouse model in multiple hematopoietic tissues.

For the SGM3 cohorts injected with IDUA-edited HSPCs, cells from the bone marrow were recovered to set up an *ex vivo* assay to assess IDUA secretion (fig 4g). Briefly, bone marrow cells were enriched for human cells and cultured *in vitro* for 48h before assessing IDUA produced intracellularly and secreted to the culture media. Lineage distribution after enrichment of animals analyzed in this assay is shown in figure 4h to aid results interpretation. We detected IDUA in the cell pellets of all conditions, with averages of 15, 56, and 30 fg/harvested cell for unedited, Ctrl-PGK and *CD11b* edited cells, respectively (fig 4i). In the supernatants, we detected IDUA only for edited conditions, with averages of 7 and 4 fg/cell in Ctrl-PGK and *CD11b* conditions, respectively (fig 4j). Of note, in *CD11b* edited conditions the main exogenous source of IDUA in cell pellets and secreted was presumably the myeloid compartment, which only accounted for 20 to 48% of the human cells seeded (fig 4h). These observations indicate that engrafted *CD11b* edited cells preserved the capacity to express and secrete supraphysiological levels of IDUA.

### Edited HSPCs as a vehicle for the delivery of therapeutics across the blood brain barrier

One of the biggest challenges in treating inborn metabolic diseases using enzyme cross-complementation is the delivery of the missing enzyme to the brain. HSPC gene therapy could be indicated for this application, because cells derived from HSPCs can cross the blood brain barrier and establish in the brain^19^. We thus evaluated whether this was true for *CD11b* edited HSPCs in our *in vivo* cohorts. We first assessed the presence of human cells (non-edited and edited) in the brains of animals injected with HSPCs edited with the eGFP or IDUA transgenes. Human cells were able to engraft in the brain tissue in both mouse models evaluated but did so with greater efficiency in SGM3 mice than in NSG mice (average of 15.1% vs. 1.0%, Fig 5a). This discrepancy could be due to the systemic expression of additional myeloid cytokines in the SGM3 model, and has been observed in previous publications^20^. No differences in brain chimerism were observed between animals injected with unedited or edited HSPCs for any mouse model (Fig 5a). We then evaluated the proportion of human cells expressing *CD11b* in brain, considered a popular microglial marker (Fig 5b) ^21^. In the brain of NSG animals injected with *CD11b* edited HSPC, where the average human fraction accounted for 0.6% of cells, the proportion of *CD11b*+ cells was significantly lower compared with the unedited group (14% vs. 41%, respectively). However, no statistical differences were observed between the *CD11b* and Ctrl-PGK conditions.

**Figure 5.**
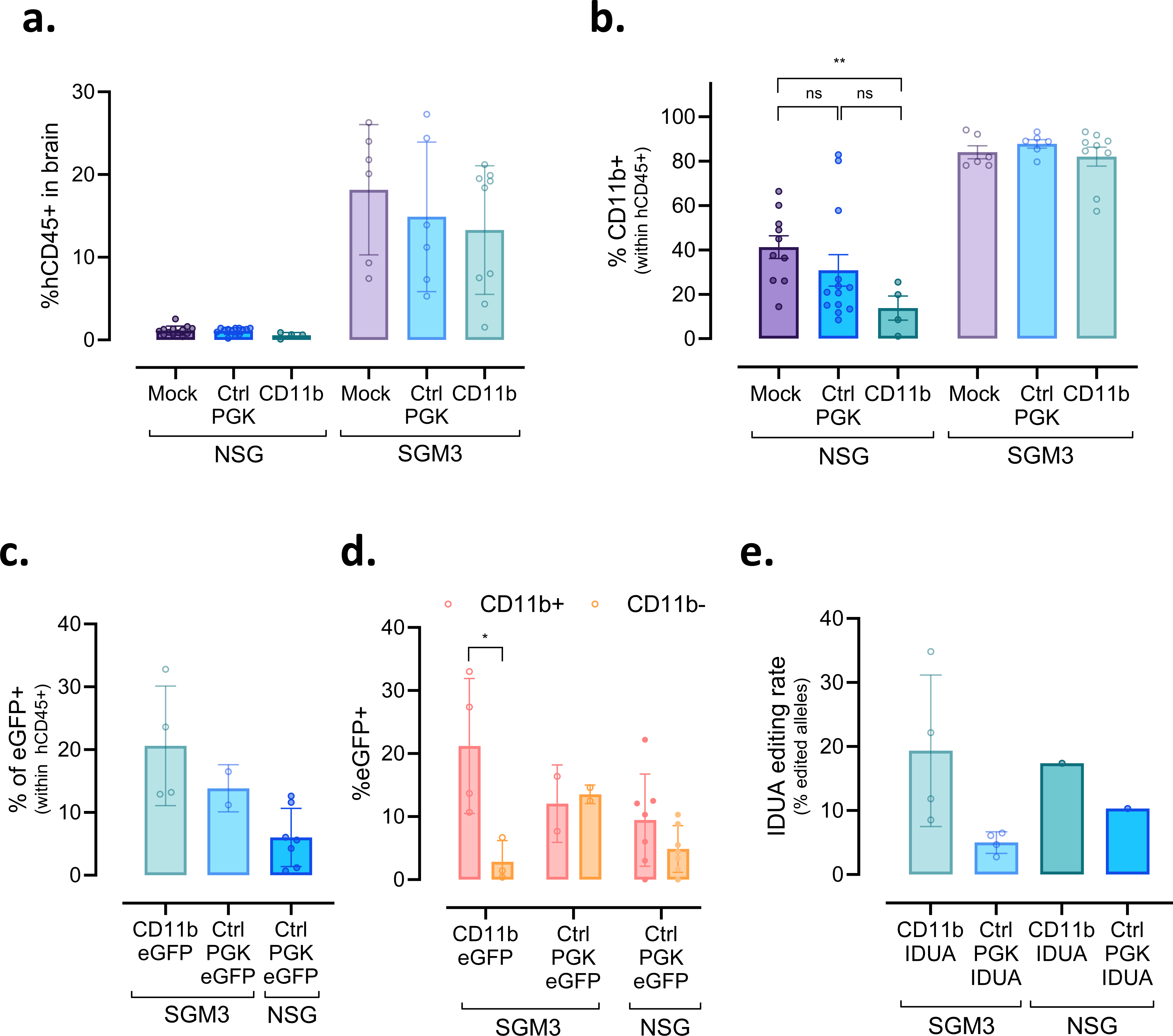
Edited HSPCs as a vehicle for therapeutic delivery across the blood-brain barrier. **a.** Engraftment in brain, measured as the proportion of human cells (hCD45+) in the leukocyte compartment in NSG and SGM3 animals injected with HSPC edited with the eGFP or IDUA cassettes. **b.** Percentage of *CD11b*+ among the human cells from animals injected with HSPC edited with the eGFP or IDUA cassettes. **c.** Percentage of eGFP+ cells within the human fraction of cell isolated from the brain of animals engrafted with HSPC edited with eGFP cassettes. **d.** Percentage of eGFP+ cells within the human *CD11b*+ and *CD11b*- fractions from the brain of animals engrafted with HSPC edited with eGFP cassettes. **e.** Percentage of edited alleles among human cells found in the brain of animals engrafted with HSPC edited with IDUA cassettes. Empty circles = SGM3. Filled circles = NSG. Data are represented as mean ± standard deviation. Statistical comparisons were performed using unpaired T tests when comparing two groups (fig 5d), or one way ANOVA when comparing three groups followed by a Tukey’s Honestly-Significant Difference post-hoc test between each two groups (fig 5a, 5b). Tukey’s p values are only shown if ANOVA was significant; ns= non-significant, *p≤0.05.

Furthermore, this difference was not observed in the SGM3 model, which displayed a much more robust human engraftment in brain for all conditions. We then characterized the presence of transgenes in human cells in the brain using flow cytometry (eGFP transgene, Fig 5c and 5d) or digital droplet PCR (IDUA transgene, Fig 5e). In SGM3 mice injected with eGFP-edited HSPC we observed an average of 20.6% and 13.9% eGFP+ cells for the *CD11b* and Ctrl-PGK groups, respectively (fig 5c), while the value for NSG mice injected with Ctrl-PGK edited cells was lower at 6.0%. Importantly, in the SGM3 cohort injected with *CD11b*-eGFP edited HSPC, we observed a myeloid-specific expression pattern, with an average of 21.2% compared to 2.8% of eGFP+ cells within the *CD11b*+ and CD11-compartments, respectively (fig 5d). Conversely, similar rates of eGFP expression were seen in both *CD11b*+ and *CD11b*- compartments for NSG and SGM3 animals injected with Ctrl-PGK-eGFP cells. This data provides additional confirmation of the ability of the *CD11b* intron editing approach to induce transgene expression specifically within the myeloid lineage of the brain tissue.

In SGM3 animals injected with IDUA edited cells, editing rates averaged 5.9% and 19.3% for the Ctrl-PGK and *CD11b* group, respectively (Fig 5e). Given the low human chimerism in the brain of NSG animals, samples from each treatment group were pooled and resulted in an editing rate of 10.3% and 17.4% for the Ctrl-PGK and *CD11b* group, respectively (Fig 5e). Together, these data confirmed that *CD11b* edited HSPCs can efficiently establish in the brain after transplantation and are able to induce the myeloid-specific expression of a given transgene.

## Discussion

The goal of this study was to develop an intron-specific gene editing approach for HSPCs that could restrict the expression of a given transgene to the myeloid lineage with a minimal genomic footprint. As a proof of concept, we targeted the endogenous *CD11b* gene given its highly specific myeloid expression pattern^22^. By design, HSPCs edited at the first intron of *CD11b* displayed transgene expression only after *in vitro* (Fig 1e) or *in vivo* (Fig 4b) myeloid differentiation, in stark contrast with the pan-lineage transgene expression pattern observed for HSPCs edited with a safe harbor approach relying on a ubiquitous promoter (Fig 1e, 4c, 4d). We then used this intron editing approach to successfully express a protein of interest, demonstrating its versatility and potential applicability for gene therapy purposes.

Targeting introns in nuclease assisted gene editing approaches has the advantage of minimizing deleterious effects that indels might have at the targeted region. Indeed, editing intronic regions to express a transgene of interest has been reported in multiple studies^23,24,25,26,27^. However, we repurposed this strategy by using a promoter-less transgene that hijacks the regulatory element of the myeloid specific *CD11b* gene without affecting its endogenous expression. We observed a seamless insertion of our cassette in the first intron of *CD11b* at the DNA level and confirmed adequate splicing yielding a bicistronic mRNA in absence of abnormal transcripts (Fig S2 and S3). This resulted in the preservation of *CD11b* endogenous expression in edited cells (Fig 1g), demonstrating the minimal footprint of our editing strategy in contrast to other gene editing approaches^28^.

We first selected eGFP as a tool to establish proof of concept, but our intron editing approach can be leveraged to express a wide variety of transgenes for therapeutic applications.

Using MPS-I as a disease model, we further validated this approach by inducing myeloid-specific expression of IDUA. Our TALEN-mediated editing at the *CD11b* intron induced up to 19-fold increase in IDUA secretion (10-fold average) by myeloid cells *in vitro* (fig 2h). We have not evaluated if these levels were sufficient to rescue the disease symptoms. However, IDUA levels obtained in our study approach those obtained by another research group (10-25 fold) which edited HSPC with a ubiquitous promoter and reported the rescue of common phenotypic impairments in an MPS-1 murine model^17^. Even lower levels of 10-16 fold induced by a strategy where an IDUA expression cassette was inserted into the albumin locus of hepatocytes were sufficient to hold therapeutic benefit in a disease mouse model^29^. Whether editing the intron of *CD11b* or perhaps another myeloid gene with stronger expression could induce therapeutic levels of IDUA or other enzymes for the treatment of LSDs warrants further studies.

It is worth noting that IDUA levels achieved are relatively high for a strategy that limits transgene expression to the myeloid lineage. For example, in another study where HSPCs were edited with a glucocerebrosidase transgene controlled by an exogenous myeloid-specific promoter for the treatment of Gaucher disease, levels of transgene overexpression by myeloid cells were only 2-fold compared to unedited cells, and 10-fold when using an ubiquitous promoter^11^.

In addition to the *CD11b* locus, this intron editing approach can be applied to a wide variety of myeloid genes displaying various levels of endogenous expression. This versatility may enable users to tailor the level of therapeutic protein secretion according to the needs of a specific pathology. In addition, other lineage- or multi-lineage-specific genes could be targeted to induce transgene expression in a specific desired compartment^30^. Safeguard systems could also be implemented into the approach to temporarily inhibit or boost gene expression using a reversible drug-inducible system^31^.

From a broader perspective, the *CD11b* intron editing strategy may represent the foundation of a powerful therapeutic platform for the treatment of inborn metabolic diseases and other neurological conditions. Indeed, myeloid-specific expression could be particularly useful for LSDs that predominantly affect the myeloid compartment like Gaucher disease, in which the accumulation of glucocerebrosides in macrophages leads to the formation of Gaucher cells^32^. In addition, lineage-specific transgene overexpression could circumvent the toxicity of certain LSD enzymes at the HSPC level. While IDUA overexpression was expected to be innocuous for HSPC^17^, some toxicity at the HSPC level has been reported after overexpression of certain LSD enzymes like galactocerebrosidase for Krabbe disease^10^ or glucocerebrosidase for Gaucher disease^11^. Further work is now needed to unfold the maximum potential of intron editing technology.

Another important advantage of using *CD11b* intron editing is to generate cells that can reach the brain compartment and deliver genetically encoded therapeutics. This is the case for microglial cells, which express substantial levels of *CD11b*^33^, especially when they are derived from HSPC after a bone marrow transplant^34^. We demonstrated that *CD11b* edited HSPCs were able to efficiently engraft in the brain of NSG and SGM3 mice and display robust transgene expression (Fig 5c). Unexpectedly, lower levels of microglial cells were observed in the *CD11b* condition compared with the Ctrl-PGK in the NSG model, but not in the SGM3 model in a context of higher brain human chimerism.

Notably, edited rates assessed in cells engrafted in the brain seem higher for the *CD11b* than the Ctrl-PGK group in this study (fig 5c, fig 5e), irrespective of the inserted cassette (GFP or IDUA) or mouse model (NSG or SGM3), even with relatively similar input *in vitro* editing rates determined before injection (fig 4a, 4e). However, previous studies using a safe harbor gene editing strategy for the overexpression of IDUA in HSPC reported strong presence of edited cells in the brain compartment, in animals that were also conditioned with busulfan^20^. Many variables could be responsible for such differences, including differences in editing rates of the subset of cells able to cross the blood brain barrier, the differentiation/expansion efficiency of edited cells after initial brain engraftment, or the role of transgene expression in such processes. While this favorable engraftment potential of *CD11b* edited HSPC remains to be confirmed in future studies, this *CD11b* intron editing approach have shown that it can assist efficient delivery of transgenes in the brain, which could be leveraged for the treatment of many metabolic and neurological diseases affecting the brain compartment, including Alzheimer’s and Parkinson’s diseases^2,35^.

One general limitation of most HSPC editing strategies based on nuclease assisted integration of a large cassette is the stability of the editing signature during the differentiation process and *in vivo* engraftment. Indeed, our data showed a decrease in editing rates *in vitro* from HSPCs to differentiated myeloid cells (Fig 1d and 2d), and *in vivo*, from injection to recovery 16 weeks after infusion (Fig 4a-4f). This is a relatively common observation that is generally attributed to the lower editing efficiency of quiescent long-term hematopoietic stem cells with respect to other CD34+ subsets^36^. This pattern could also be attributed to the well-known toxicity of the AAV viral particles used for transgene delivery^37,38^. Vectorizing the transgene using non-viral DNA delivery methods like single strand DNA electroporation may improve the fitness of edited HSPCs and promote their long-term maintenance *in vivo* as proposed earlier^38^. Further work is now needed to explore this avenue beyond the scope of this manuscript.

In summary, we describe here a TALEN-mediated intron editing approach in HSPCs that enables myeloid-specific expression of a transgene without ectopic expression in the HSPCs. This approach enables edited myeloid cells to populate multiple hematopoietic tissues and cross the blood brain barrier to engraft in the brain as microglial cells, making it attractive for delivering genetically encoded therapeutics to the brain compartment. We believe this editing approach is a valuable tool added to the HSPC gene therapy toolbox that could complement current editing strategies in the development of therapeutics for genetic and neurodegenerative conditions.

## Materials and Methods

### HSPC isolation and culture

Fresh human mobilized LeukoPaks from healthy donors were purchased from AllCells (Alameda, CA, USA). CD34+ HSPCs were isolated from the mobilized LeukoPaks using a CliniMACS Plus Instrument, CliniMACS CD34 GMP MicroBeads, and CliniMACS Tubing Set LS (Miltenyi Biotec, Bergisch Gladback, Germany). CD34+ and CD34- cell fractions were stained with CD34-PE (Clone: REA1164; Miltenyi) and CD45-APC (Clone REA747; Miltenyi). Sample purity was assessed on a NovoCyte Penteon Flow Cytometer (Agilent Technologies, Santa Clara, CA, USA). After enrichment, the CD34+ fractions were vitally frozen using CryoStor CS10 (StemCell Technologies, Vancouver, Canada).

Cells were thawed and recovered from frozen aliquots and cultured at 37°C, 5% CO2 for 48 hours prior to electroporation at a seeding concentration of 4.0 x 10^5^ cells/mL in HSPC culture media (StemSpan SFEM II (StemCell Technologies) supplemented with penicillin/streptomycin (100 U/mL; ThermoFisher Scientific, Waltham, MA, USA) and 1X CD34+ Expansion supplement (StemCell Technologies).

### TALEN and mRNA Production

TALEN pair used for editing the *CD11b* locus targeted 5’ – ACAACATATTCTATC – 3’ and 5’ AGTAAATTTTAGGTTT – 3’ sequences, located between its first and second exon. TALEN pair used for editing the AAVS1 locus targeted 5’ – CCCCTCCACCCCACA – 3’ and 5’ – GTCACCAATCCTGTC – 3’ (Fig S6a). mRNA for most experiments was purchased from TriLink BioTechnologies (San Diego, CA, USA). For AAVS1 TALEN mRNA, some in house productions were generated using the T7-FlashScribe Transcription Kit combined with the ScriptCap m^7^G Capping System and the A-Plus Poly(A) Polymerase Tailing Kit (CellScript, Madison, WI, USA) according to the manufacturers’ protocols. mRNA quality was assessed via Fragment Analyzer (Agilent Technologies).

### AAV donor plasmid construction and production

AAV6 donor sequences used to edit the *CD11b* and AAVS1 loci with eGFP or IDUA inserts (Fig S6b) were designed using Geneious software and ordered from GenScript (Piscataway, NJ, USA). Nucleotide sequences are depicted in Table S1. The constructs were assembled into the AAV2-ITR plasmid via molecular cloning and purified using the NucleoBond Xtra Maxi EF kit (Macherey-Nagel, Düren, Germany). The purified plasmids were sent to Vigene Biosciences (now Charles River, Wilmington, MA, USA) VectorBuilder (Chicago, IL, USA) to produce AAV6.

### HSPC gene editing

CD34+ HSPCs were transfected 48 hours after thawing and expansion by combining 5 μg of each AAVS1 TALEN mRNA or 10 μg of each *CD11b* TALEN mRNA with 1e6 HSPC in 100 μL BTXpress Solution (BTX Technologies, Hawthorne, NY, USA) and electroporating using the PulseAgile (Cellectis, Paris, France). One μg of Via-Enh01 mRNA and 4 μg of HDR-Enh01 mRNA were also included in the electroporation mix because Via-Enh01^39^ protein inhibits cell apoptosis and because HDR-Enh01^40^ increases homologous recombination by inhibiting the non-homologous-end-joining pathway. Following electroporation, cells were immediately mixed with HSPC culture media. Cells were seeded in a 96-well plate at a concentration of 2.0 x 10^6^ cells/mL and transduced with AAV serotype 6 (MOI 1.0 x 10^4^ vg/cell). For editing with a safe harbor approach, the AAV sequence comprised a PGK promoter followed by eGFP or codon optimized IDUA sequences flanked by homology sequences for the AAVS1 locus. For editing the first intron of *CD11b*, the AAV sequence comprised 5’ and 3 splicing sequences flanking an eGFP or codon optimized IDUA sequences followed by a 2A viral peptide sequence and a reencoded first exon of the *CD11b* locus. This construct was flanked by homology arms for the *CD11b* gene. Nucleotide sequences are described in Table S1. A mock-electroporated control and TALEN-only controls were included in most experiments. After transfection/transduction, edited HSPCs were incubated at 37°C for 15 minutes and transferred to 30°C overnight (16-20h) to then be cultured at 37°C for the remaining follow-up.

### Measuring insertions at the AAVS1 and *CD11b* loci using digital droplet PCR

Cell pellets were harvested from bulk CD34+ HSPCs four to five days post-editing, from bulk myeloid differentiated cells 13-15 days post-editing, or from cellular suspensions of multiple mouse tissues (blood, bone marrow, spleen, or brain). Genomic DNA was extracted using the Mag-Bind Blood & Tissue DNA HDQ 96 Kit (Omega Bio-Tek, Norcross, GA, USA) according to the manufacturer’s protocols on the KingFisher Flex purification system (ThermoFisher Scientific). A multiplex digital droplet PCR (ddPCR) assay was performed to measure gene editing rates, where one primer/probe set was designed to amplify a reference gene (*CCR5*) and another set amplified the integrated cassette at the desired genomic location (AAVS1 or *CD11b*) using an in-out PCR approach. For *CCR5*, primers sequences were 5’-AAATAAGCTGCCTTGAGCC-3’ and 5’-TGTTGCACTCTCCACAACTT-3’, and a HEX-tagged 5’-TCCCTTCGTTGCTTCCTGCTGACA-3’ probe. For AAVS1, primers sequences were 5’- GTAGTCTGGCACTGGAGC-3’ and 5’-GACGGATGTCTCCCTTGC-3’, and a FAM-tagged 5’-CCTCCCCTTCTTGTAGGCCTGC-3’ probe. For *CD11b*, primers sequences were 5’- GGCTCTCAGAGTCCTTCTGT-3’ and 5’-TCACTTGAGCCCTGATTGTG-3’, and a FAM- tagged 5’-TCCAGCCTGAGCAACAGAGA-3’ probe. A total of 60-100 ng gDNA was combined with 22.5 pmol each forward and reverse target primers, 22.5 pmol each forward and reverse reference primers, 62.5 pmol each of target FAM probe and reference HEX probe, 1X ddPCR Supermix for Probes without dUTP (Bio-Rad Laboratories, Hercules, CA, USA), and 10 units of XbaI (New England Biolabs, Ipswich, MA, USA) in a 25 µL reaction. Droplets were generated on a QX200 Droplet Generator (Bio-Rad Laboratories) according to the manufacturer’s protocol. Droplets were amplified using a Bio-Rad PCR thermocycler using the following PCR conditions: 95°C for 10 min, 40 cycles of 94°C for 30 sec and 60°C for 2 min 30 sec, followed by 98°C for 10 min and 4°C until droplet analysis. Droplets were analyzed on a QX200 Droplet Reader (Bio-Rad Laboratories) using the QuantaSoft Software (Bio-Rad Laboratories) to detect FAM and HEX fluorescence positive and negative droplets according to the manufacturer’s protocol. Control samples with non-template control and mock-treated samples were included.

### *In vitro* myeloid differentiation

Twenty-four hours after gene editing, approximately 40,000 non-adherent HSPCs were transferred to new 96-well polystyrene plates and cultured with 200 μL of myeloid II differentiation medium, containing StemSpan SFEM II (StemCell Technologies) supplemented with penicillin/streptomycin (100 U/mL; ThermoFisher Scientific) and 1X Myeloid Expansion Supplement II (StemCell Technologies). Cells were maintained for up to three weeks in the same plates and split every three to four days by discarding 160 μL of medium and adding 160 μL of fresh differentiation medium to each well.

### Colony forming unit assays and clonal genotyping

Mock-treated, Ctrl-PGK edited, and *CD11b* edited CD34+ HSPCs were counted and plated in duplicates in a SmartDish™ (StemCell Technologies) at a density of 200 cells/well in 1.1 mL of methylcellulose (Methocult H4435 Enriched, StemCell Technologies). This media contains SCF, IL3, IL6, erythropoietin, G-CSF, and GM-CSF, which support the growth of blood progenitor cells, including erythroid progenitors (burst forming unit-erythroid or BFU-E, and colony-forming unit erythroid or CFU-E), granulocyte-macrophage progenitors (CFU-GM), and multi-potential granulocyte, erythroid, macrophage, megakaryocyte progenitor cells (CFU-GEMM). After 14 days, the colonies were imaged, counted, and classified using the STEMvision (StemCell Technologies) according to the ‘STEMvision™ Automated Colony-Forming Unit (CFU) Assay Reader’ manual.

### IDUA secretion assays and ELISA

CD34+ HSPCs or myeloid differentiated cells were seeded at a density of 5.0 x 10^5^ cells/mL in untreated 96-well polystyrene plates in HSPC culture media and cultured at 37°C, 5% CO2. After 72 h, the non-adherent cells were harvested from the plates, resuspended in 200 μL of PBS, and lysed by 3 rounds of freeze-thaw cycles at -80°C. The supernatants were also collected and stored at -20°C. After lysis, the remaining cellular debris was removed by centrifugation at 10,000 x *g* for 15 min, and the supernatants were recovered.

IDUA enzyme expression was measured fluorometrically in a 96-well plate using a Human IDUA ELISA kit (Sigma-Aldrich, St. Louis, MO, USA) according to the manufacturer’s sandwich assay protocol. Standards and samples were incubated in the provided plate overnight at 4°C. At the end of the procedure, the plate was read at 450 nm with the FLUOstar Omega Plate Reader (BMG LabTech, USA). IDUA activity was assessed using 4-methylumberlliferyl-alpha-L-iduronide (Glycosynth, Cheshire, England) and recombinant human alpha-L-iduronidase protein (R&D systems, Minneapolis) following manufacturer’s instructions.

### Flow cytometry

For analysis of eGFP integration and cell phenotype, 50,000 to 200,000 cells were harvested per condition, washed in staining buffer (PBS containing 1mM EDTA and 0.5% BSA), and resuspended in 0.25 ug of Human Fc Block (BD Biosciences, Franklin Lakes, NJ, USA) diluted in staining buffer to block non-specific antibody binding. Cells were stained (30 min, 4°C in the dark) with *CD11b* (Clone: REA713; Miltenyi or Clone: M1/70; Biolegend, San Diego, CA, USA), CD14 (Clone: TUK4 or Clone: REA599; Miltenyi), CD15 (Clone: W6D3; Biolegend), CD33 (Clone: REA775; Miltenyi), and CD34 (Clone: 581; Biolegend). Cells were then washed with staining buffer, fixed, and permeabilized (20 min, 4°C, dark) using Fixation and Permeabilization Solution (BD Biosciences). Cells were then stained with S100A9 (Clone: MAC387; Invitrogen, Waltham, MA, USA) (30 min, 4°C in the dark). Fluorescence was detected and analyzed on a NovoCyte Penteon Flow Cytometer (Agilent Technologies).

### Mic

NOD.Cg-Prkdc^scid^Il2rg^tm1Wjl^/SzJ (NSG) and NOD.Cg-Prkdc^scid^Il2rg^tm1Wjl^Tg(CMV- IL3,CSF2,KITLG)1Eav/MloySzJ (NSG-SGM3) mice were developed at and ordered from The Jackson Laboratory (Bar Harbor, ME, USA). The mice were housed at a Mispro Biotech Services (New York, New York) in a room with a 12-h light/dark cycle that was temperature- and humidity-controlled. The mice were given unlimited access to sterile food and water. All experiments were performed in accordance with the National Institutes of Health’s institutional guidelines and were approved by the Institutional Animal Care and Use Committee (IACUC).

### Transplantation of human CD34+ HSPCs into NSG or SGM3 mice

Mice were conditioned with busulfan via intraperitoneal injections at 15mg/kg of busulfan for one (SGM3) or three consecutive days (NSG) before HSPC transplantation. A total of 0.75 x 10^6^ mock-electroporated or edited CD34^+^ HSPCs were retro-orbitally injected into 6-8-week-old female mice 24-hours post-editing. Human engraftment was evaluated in peripheral blood harvested via retro-orbital bleeding at 4, 8, and 12 weeks post-transplantation. At 15-18 weeks post-transplantation, mice were euthanized, and peripheral blood (PB), spleen, bone marrow (BM), and brain were harvested and analyzed for human engraftment and IDUA expression. All procedures involving animals were approved by The Mispro Institutional Animal Care and Use Committee (IACUC) and were performed in accordance with the guidelines of the Public Health Service (PHS) Policy on Humane Care and Use of Laboratory Animals, Office of Laboratory Animal Welfare (OLAW), and the United States Department of Agriculture (USDA) Animal Welfare Act (AWA).

### Necropsies and tissue analysis

At 15-18 weeks post-transplantation, mice were sedated with isoflurane, peripheral blood was extracted via cardiac puncture, and transcardial perfusion was performed with ice-cold PBS. Mice were decapitated, and brains were collected in ice-cold Dulbecco’s Modified Eagle Medium (DMEM; ThermoFisher Scientific) mixed with penicillin/streptomycin (100 U/mL; ThermoFisher Scientific). The spleens, femur, and tibia of both legs were extracted into ice-cold PBS. All organs were kept on ice until ready for processing.

Briefly, BM cells were collected via centrifugation of cut femurs and tibiae; splenocytes were collected by mashing the spleens on a 45μm mesh strainer; and neurons and neuroglia were collected by homogenizing the brain with a 16-gauge needle, followed by enzymatic dissociation with a 1:1 mixture of papain and DNase I, followed by Percoll density gradient centrifugation. The mononuclear cells were recovered, non-specific antibody staining was blocked with Human Fc Block (BD Biosciences) for 5 min at 4°C, and the cells were stained at 4°C for 30 min in the dark. The following antibody panel was used to assess human engraftment in cells harvested from bone marrow and spleen from eGFP-edited mouse cohorts and from peripheral blood in both eGFP- and IDUA-edited mouse cohorts: hCD45-APC (Clone: REA747; Miltenyi), mCD45-V450 (Clone 30- F11; BD Biosciences), CD3-PerCP-Cy5.5 (Clone: UCHT1; BD Biosciences), CD33-PE (Clone: AC104.3E3; Miltenyi), CD34-APC-Vio770 (Clone: REA1164; Miltenyi), and CD19-PE-Vio770 (Clone: 555412; BD Biosciences). For neurons and neuroglia from both eGFP-edited and IDUA- edited mouse cohorts, the following antibody panel was used: hCD45-PE-Cy7 (Clone: REA747; Miltenyi), mCD45-APC-Cy7 (Clone: 30-F11; BD Biosciences), and CD11b-PE (Clone: M1/70; Biolegend). For IDUA-edited mouse cohorts, the following antibody panel was used to assess human engraftment in cells harvested from the bone marrow and spleen: hCD45-PE-Cy7 (Clone: REA747; Miltenyi), mCD45-V450 (Clone: 30-F11; BD Biosciences), CD19-FITC (Clone: 555412; BD Biosciences), CD3-PerCP-Cy5.5 (Clone: UCHT1; BD Biosciences), CD33-APC (Clone: AC104.3E3; Miltenyi), CD34-APC-Vio770 (Clone: REA1164; Miltenyi), and CD11b-PE (Clone: M1/70; Biolegend).

### Characterization of cassette insertion and RNA splicing

Genomic DNA was extracted from IDUA-edited cells after 14 days of myeloid differentiation as described above. From paired samples, total RNA from mock-electroporated and IDUA-edited cells was isolated using TRIzol (Invitrogen) according to the manufacturer’s protocol. cDNA was synthesized using SuperScript III (Invitrogen) and Oligo(dT) primers (Invitrogen) per the manufacturer’s protocol. RNase H (ThermoFisher Scientific) was used to remove RNA template from the cDNA following the manufacturer’s protocol. *CD11b*-specific amplicons were generated via 3-step nested PCR using PrimeSTAR GXL (Takara Bio, Kusatsu, Shiga, Japan) according to the manufacturer’s protocol with a 60°C annealing temperature, 6.5 min extension, and a 35-cycle reaction. The first PCR was set up with a forward primer inside the first exon (5’- GTGGTTCCTCAGTGGTGCCTGCAACCC-3’) and a reverse primer inside the second exon (5’- CACATCGCCTGGAATCTGTCTGAAGG-3’). The second PCR was set up with a forward primer downstream of the first forward primer (5’-GTTCACCTCCTTCCAGGTTCTGGC-3’) and a reverse primer upstream of the first reverse primer (5’- CATCAGCGCTGGTGTGGAGGAGGTTCC-3’). The same primers were used to generate amplicons from genomic DNA in paired DNA samples. PCR products from genomic DNA and cDNA were purified using NucleoSpin Gel and PCR Clean-up Mini Kit (Macherey-Nagel, Allentown, PA, USA), and quality was validated via agarose gel electrophoresis.

DNA libraries were prepared using the SQK-LSK109 protocol and kit from Oxford Nanopore Technologies (ONT) (Oxford, UK). Libraries were barcoded with ONT’s EXP-NBD104 kit and sequenced for 72-hours on the FLO-MIN106 flow cell using the MinION platform.

The resulting reads were trimmed (approximately 50bp) to ensure reading quality and aligned on the human genome (release hg38) using minimap2. The reads aligning on *CD11b* were then aligned on the edited *CD11b* sequence. Reads with at least 100 bases aligned on the cassette (not necessarily consecutive) were considered edited. For unedited sequences of unedited or edited samples, incomplete reads due to incomplete PCR or incomplete sequencing were manually removed after confirming missing portions at the sequence initiation or end. Unedited sequences from edited samples were removed to simplify the analysis, as they were no longer representative of frequency given the amplification bias between unedited (short) and edited (long) amplicons. Finally, 300 reads were randomly selected of the expected size for non-edited (4kb) or edited (6kb) transcripts in the control and edited samples, respectively. These reads were manually confirmed to be representative, and their alignments were plotted using a custom script.

### Statistical Analysis

Data are represented as mean ± standard deviation in most graphs. Statistical comparisons were performed using unpaired T tests when comparing two groups, or one way ANOVA when comparing three or more groups followed by a Tukey’s Honestly-Significant Difference post-hoc test between each two groups. The significance threshold for statistical analyses was set to 0.05: *≤0.05; **≤0.01; ***≤0.001; ****≤0.0001; “ns” means not significant. All statistical analyses were performed using GraphPad Prism v.9.4 (GraphPad Software, Inc., San Diego, CA).

## Acknowledgments

TALEN® is a Cellectis patented technology. Data described in this paper is available upon request.

## Author contributions

E.S., J.C.J., A.O.L., A.B., and D.T. performed the experiments, S.P. analyzed nanopore data; E.S., A.B., A.J., P.D., and J.V. designed the study, and E.S. and J.V. wrote the manuscript. Figures 1a, 2a, 3a, and 4g were created using biorender.com.

