## Supplementary figures and images for "TALEN-mediated intron editing of HSPCs enables transgene expression restricted to the myeloid lineage"

### Supplemental figures S1-S6

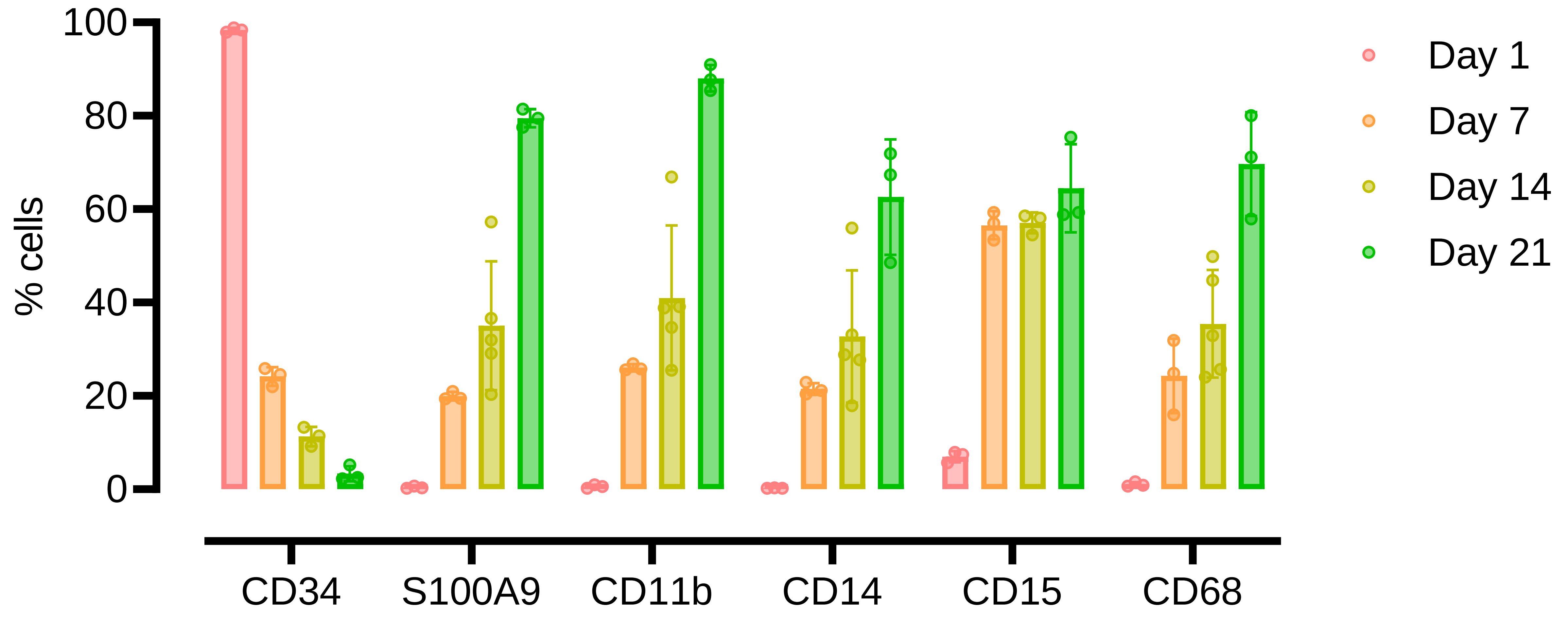

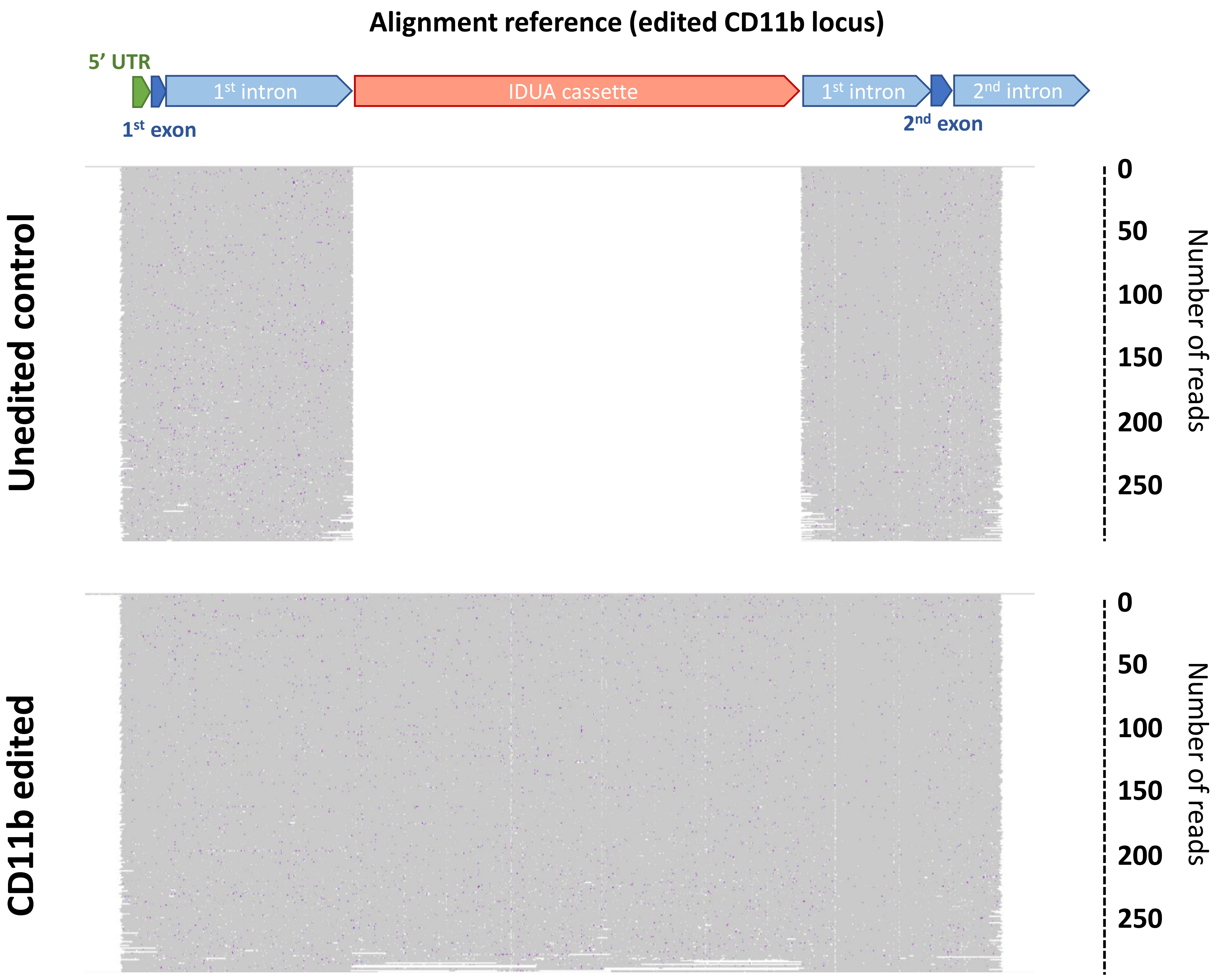

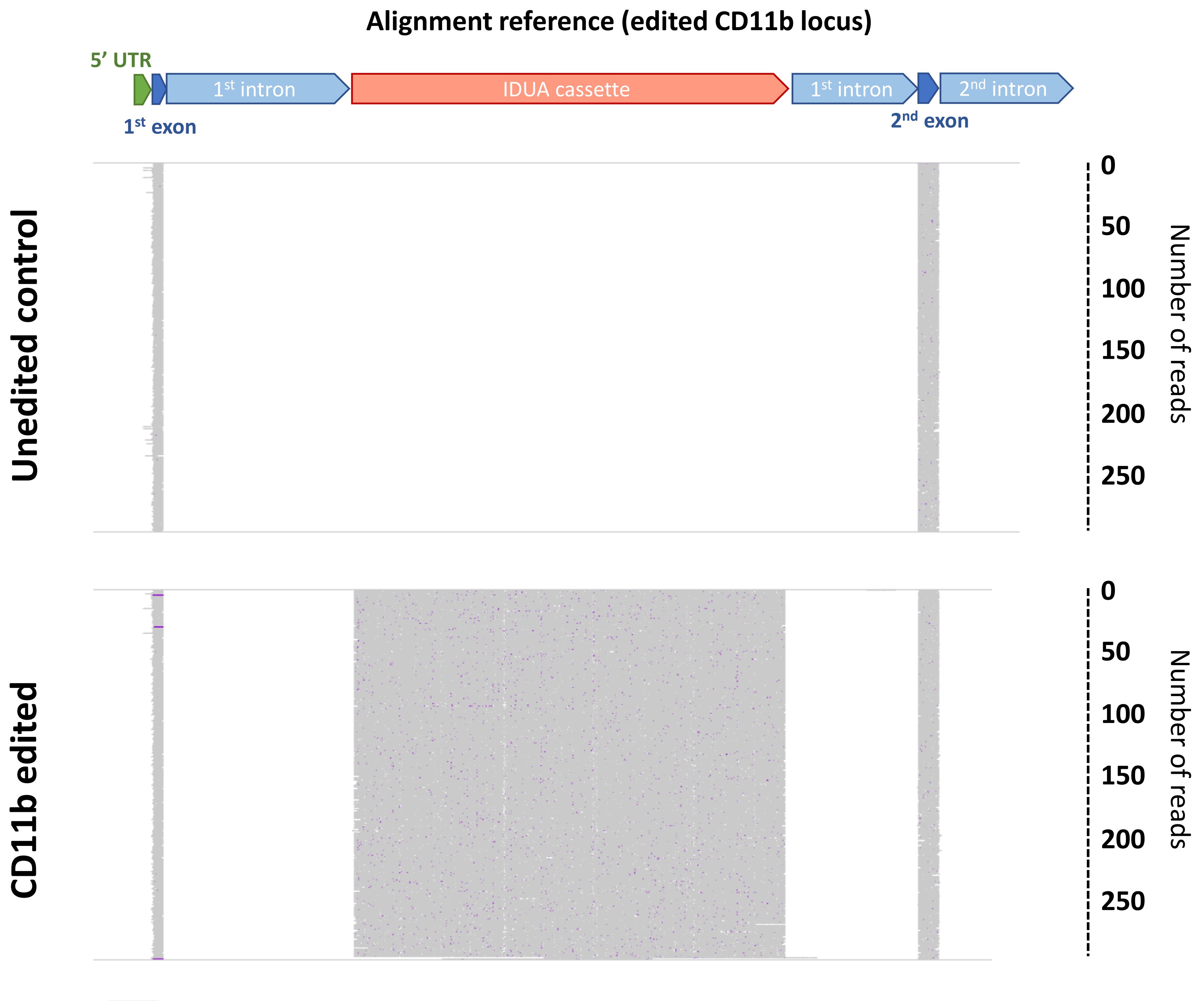

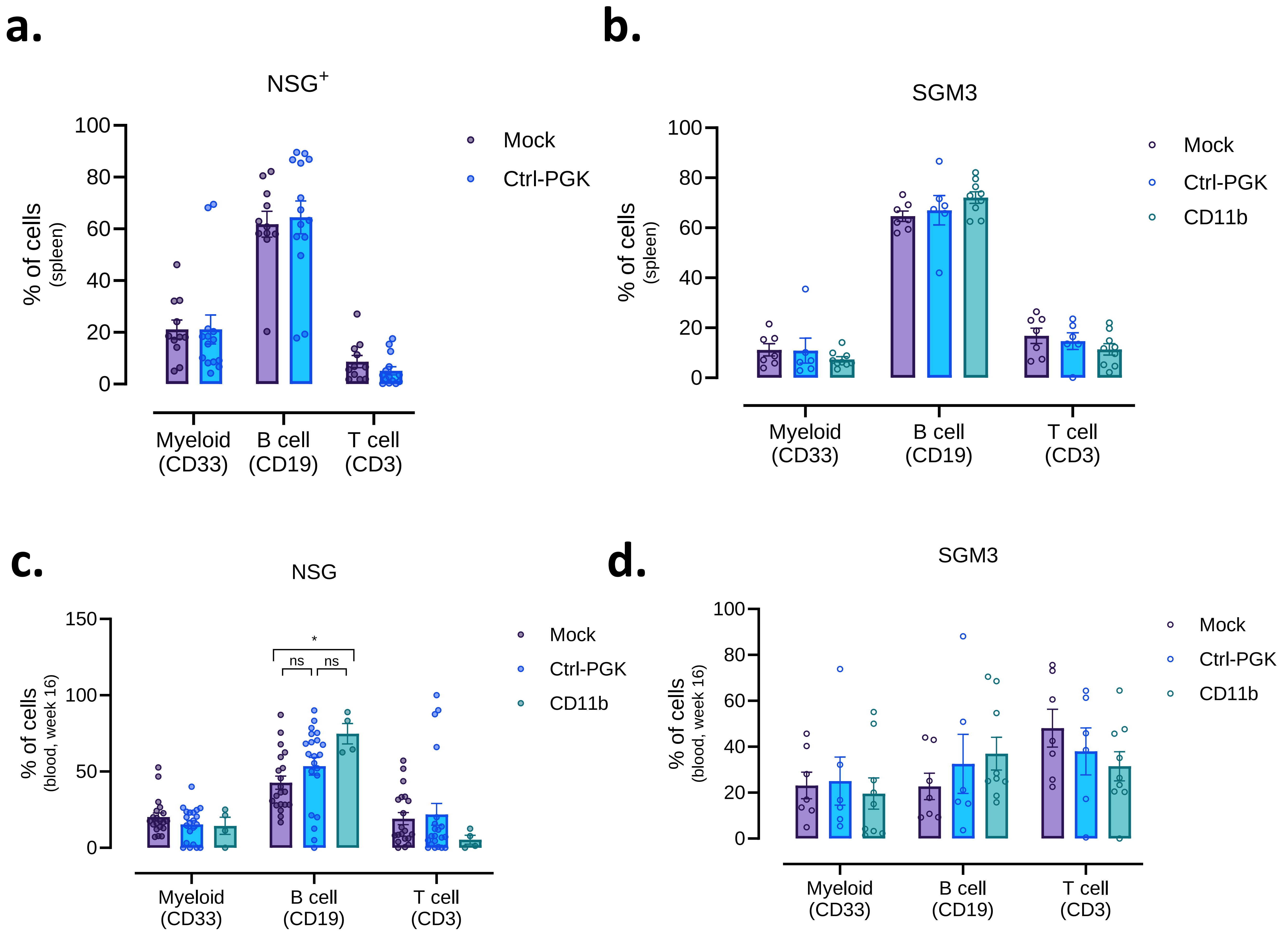

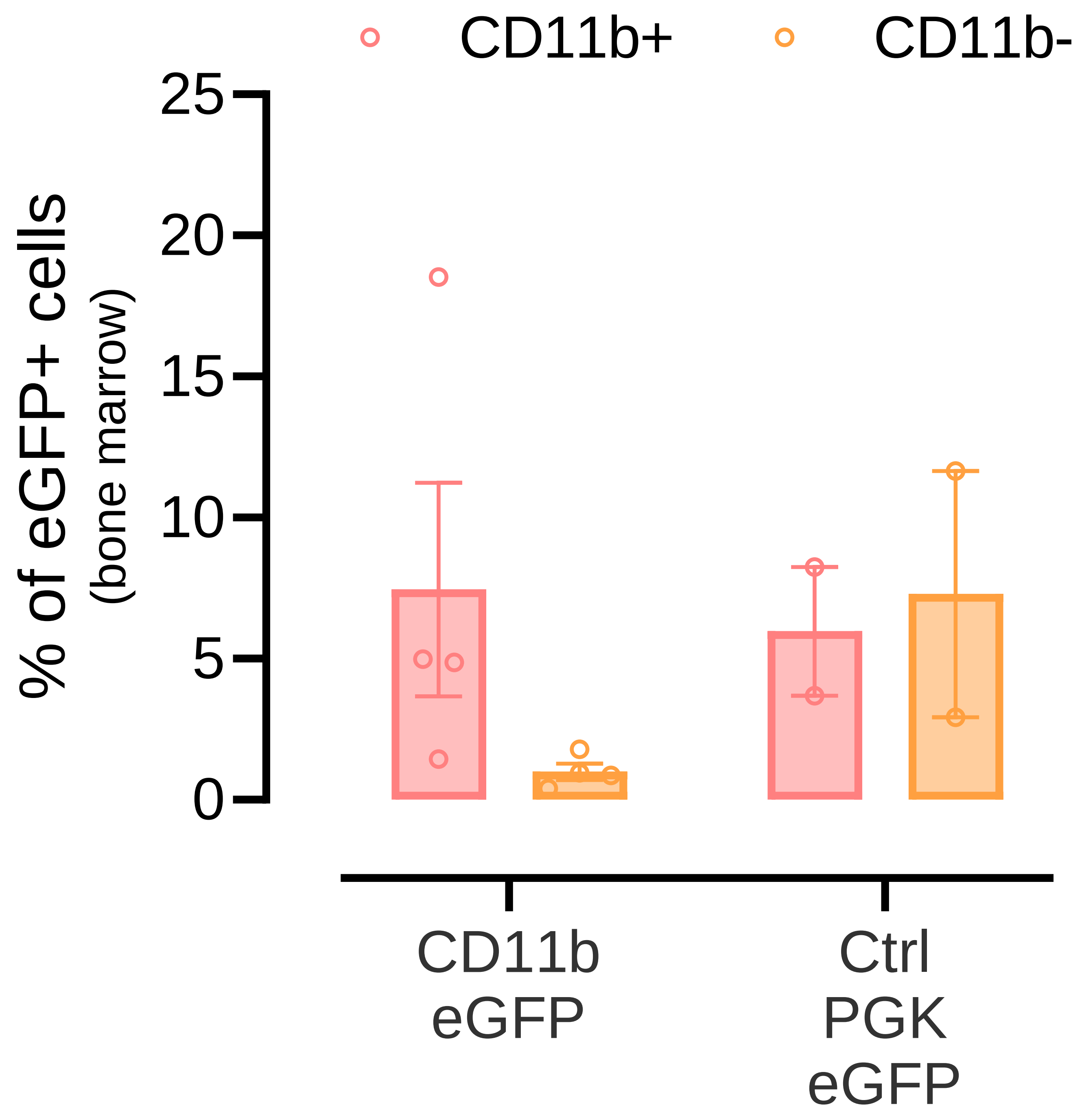

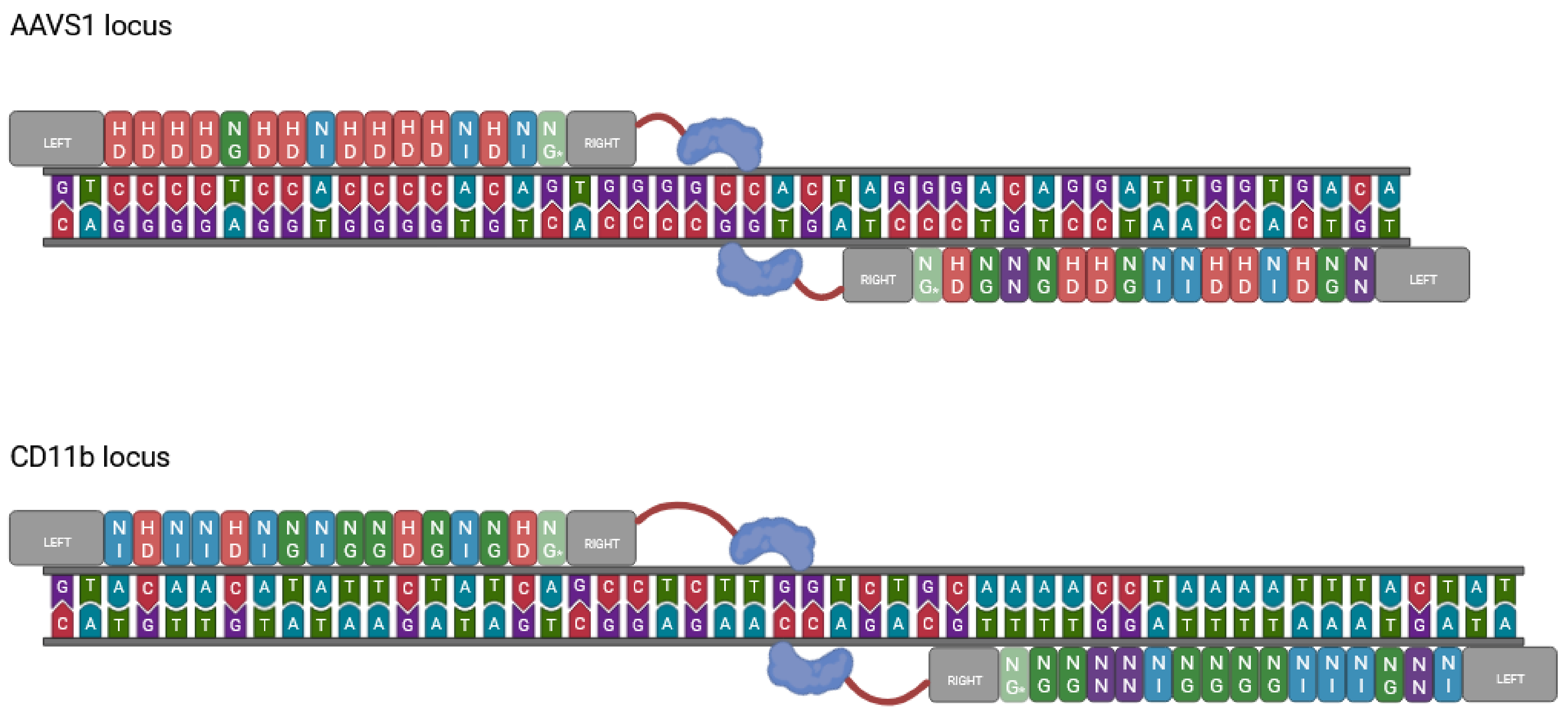

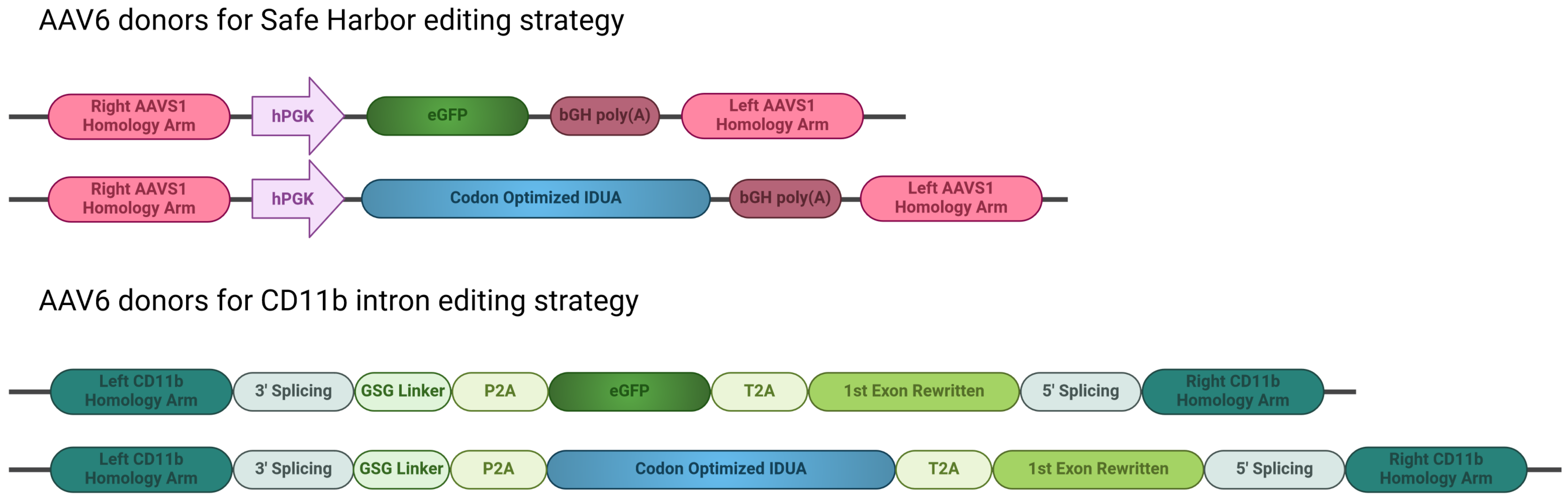
