## Supplemental figure legends and table S1 for "TALEN-mediated intron editing of HSPCs enables transgene expression restricted to the myeloid lineage"

**Figure S1. *In vitro* myeloid differentiation dynamics.** Gradual appearance of myeloid markers over time in human HSPCs with a myeloid cytokine cocktail containing fms-like tyrosine kinase 2 ligand (Flt3L), stem cell factor (SCF), thrombopoietin (TPO), macrophage colony stimulating factor (M-CSF), and granulocyte/macrophage colony stimulating factor (GM-CSF). Data are represented as mean ± standard deviation.

**Figure S2. Analysis of transgene cassette integration at the DNA level.** Alignment of gDNA sequenced by long-read sequencing (Nanopore, Oxford Technologies) from unedited and edited cells to the reference gene of a perfectly edited locus. For the *CD11b* edited sample, unedited sequences were excluded (details in methods section). Three hundred representative sequences are shown per sample; grey indicates matching base pair, purple indicates a base pair mismatch, and white indicates a missing nucleotide.

**Figure S3. Characterization of splicing in myeloid-differentiated edited cells via RNA sequencing.** Alignment of cDNA sequenced by long-read sequencing (Nanopore, Oxford Technologies) from unedited and edited cells to the reference gene of a perfectly edited locus at the DNA level. For the *CD11b* edited sample, unedited sequences were excluded (please refer to methods). Three hundred representative sequences are shown per sample; grey indicates matching base pair, purple indicates a base pair mismatch, and white indicates a missing nucleotide.

**Figure S4. Gene editing does not affect the differentiation capacity of edited HSPC.** Lineage distribution in spleen cells is represented for NSG (**a**) and SGM3 (**b**) mice 16 weeks post HSPC injection. Lineage distribution in blood cells is represented for NSG (**c**) and SGM3 (**d**) mice 16 weeks post HSPC injection. Empty circles = SGM3. Filled circles = NSG. Data are represented as mean ± standard deviation. $^{+}$No spleens were harvested for the *CD11b* group. Statistical comparisons were performed using unpaired T tests when comparing two groups (Fig S4a), or one way ANOVA when comparing three groups followed by a Tukey's Honestly-Significant Difference post-hoc test between each two groups (Fig S4b, S4c, and S4d). Tukey's p values are only shown if ANOVA was significant; ns= non-significant, *$p \leq 0.05$.

**Figure S5. HSPCs edited at the *CD11b* intron support myeloid specific transgene expression *in vivo*.** Quantification of eGFP+ cells based on *CD11b* expression from a subset of SGM3 animals injected with Ctrl-PGK or *CD11b* edited HSPC with the eGFP cassette. All animals shown are SGM3. Data are represented as mean ± standard deviation.

**Figure S6. Schematics of editing tools**. **a**. Schematics of TALEN pairs targeting the AAVS1 (top) and *CD11b* (bottom) locus, indicating binding and targeted sequence. **b**. Schematics of AAV6 donors used for editing the AAVS1 and *CD11b* loci with eGFP and IDUA cassettes. Specific sequences are depicted in Table S1.

**Table S1**. Nucleotide sequences of components from AAV6 donors used to edit AAVS1 and *CD11b* locus, as described in Fig S6b.

| | |
|---|---|
| **AAVS1 right homology arm (inverted)** | CCCACAGTTGGAGGAGAATCCACCCAAAAGGCAGCCTGGTAGACAGGGCTGG GGTGGCCTCTCGTGGGGTCCAGGCCAAGTAGGTGGCCTGGGGCCTCTGGGGG ATGCAGGGGAAGGGGGATGCAGGGGAACGGGGATGCAGGGGAACGGGGCTC AGTCTGAAGAGCAGAGCCAGGAACCCCTGTAGGGAAGGGGCAGGAGAGCCA GGGGCATGAGATGGTGGACGAGGAAGGGGGACAGGGAAGCCTGAGCGCCTC TCCTGGGCTTGCCAAGGACTCAAACCCAGAAGCCCAGAGCAGGGCCTTAGGG AAGCGGGACCCTGCTCTGGGCGGAGGAATATGTCCCAGATAGCACTGGGGAC TCTTTAAGGAAAGAAGGATGGAGAAAGAGAAAGGGAGTAGAGGCGGCCACG ACCTGGTGAACACCTAGGACGCACCATTCTCACAAAGGGAGTTTTCCACACG GACACCCCCCTCCTCACCACAGCCCTGCCAGGACGGGGCTGGCTACTGGCCTT ATCTCACAGGTAAAACTGACGCACGGAGGAACAATATAAATTGGGGACTAGA AAGGTGAAGAGCCAAAGTTAGAACTCAGGACCAACTTATTCTGATTTTGTTTT TCCAAACTGCTTCTCCTCTTGGGAAGTGTAAGGAAGCTGCAGCACCAGGATC AGTGAAACGCACCAGACGGCCGCGTCAGAGCAGCTCAGGTTCTGGGAGAGG GTAGCGCAGGGTGGCCACTGAGAACCGGGCAGGTCACGCATCCCCCCCTTCC CTCCCACCCCCTGCCAAGCTCTCCCTCCCAGGATCCTCTCTGGCTCCATCGTA AGCAAACCTTAGAGGTTCTGGCAAGGAGAGAGATGGCTCCAGGAAATGGGG GTGTGTCACCAGATAAGGAATCTGCCTAACAGGAGGTGGGGGTTAGACCCAA TATCAGGAGACTAGGAAGGAGGAGGCCTAAGGATGGGGCTTTTCTGTCACCA ATCCTGTCCCTAGT |
| **hPGK Promoter** | GGGGTTGGGGTTGCGCCTTTTCCAAGGCAGCCCTGGGTTTGCGCAGGGACGC GGCTGCTCTGGGCGTGGTTCCGGGAAACGCAGCGGCGCCGACCCTGGGTCTC GCACATTCTTCACGTCCGTTCGCAGCGTCACCCGGATCTTCGCCGCTACCCTT GTGGGCCCCCCGGCGACGCTTCCTGCTCCGCCCCTAAGTCGGGAAGGTTCCTT GCGGTTCGCGGCGTGCCGGACGTGACAAACGGAAGCCGCACGTCTCACTAGT ACCCTCGCAGACGGACAGCGCCAGGGAGCAATGGCAGCGCGCCGACCGCGA TGGGCTGTGGCCAATAGCGGCTGCTCAGCAGGGCGCGCCGAGAGCAGCGGCC GGGAAGGGGCGGTGCGGGAGGCGGGGTGTGGGGCGGTAGTGTGGGCCCTGT TCCTGCCCGCGCGGTGTTCCGCATTCTGCAAGCCTCCGGAGCGCACGTCGGCA GTCGGCTCCCTCGTTGACCGAATCACCGACCTCTCTCCCCAG |

| | |
|---|---|
| **Codon optimized IDUA** | ATGCGGCCCCTGAGGCCTAGAGCTGCCCTGCTGGCTCTGCTGGCCTCTCTGCT<br>GGCTGCCCCTCCTGTGGCCCCTGCCGAAGCCCCTCACCTGGTGCATGTGGATG<br>CCGCCAGGGCTCTGTGGCCACTGCGGAGATTCTGGCGGAGCACCGGCTTTTGC<br>CCCCCACTGCCTCACAGCCAGGCCGACCAGTACGTGCTGAGCTGGGACCAGC<br>AGCTGAACCTGGCCTACGTCGGCGCCGTGCCCCACAGAGGCATCAAACAAGT<br>GCGCACCCACTGGCTGCTGGAACTGGTGACAACCCGGGGCAGCACCGGCAGA<br>GGACTGAGCTACAACTTCACCCACCTGGACGGCTACCTGGACCTGCTGAGAG<br>AGAACCAGCTGCTGCCCGGCTTCGAGCTGATGGGCAGCGCCAGCGGCCACTT<br>CACCGACTTCGAGGACAAGCAGCAGGTGTTCGAGTGGAAGGACCTGGTGTCC<br>AGCCTGGCCAGACGGTACATCGGCAGATACGGCCTGGCCCACGTGTCCAAGT<br>GGAACTTCGAGACATGGAACGAGCCCGACCACCACGACTTCGACAACGTGTC<br>AATGACCATGCAGGGCTTTCTGAACTACTACGACGCCTGCAGCGAGGGCCTG<br>AGAGCCGCCTCTCCTGCCCTGAGACTGGGCGGACCCGGCGATAGCTTCCACA<br>CCCCCCCCAGAAGCCCCCTGAGCTGGGGCCTGCTGAGACACTGCCACGACGG<br>CACCAATTTCTTCACCGGCGAGGCCGGCGTGCGGCTGGACTACATCAGCCTGC<br>ACCGGAAGGGCGCCAGAAGCAGCATCAGCATCCTGGAACAGGAAAAGGTCG<br>TCGCCCAGCAGATCCGGCAGCTGTTCCCCAAGTTCGCCGACACCCCCATCTAC<br>AACGACGAGGCCGACCCCCTGGTCGGATGGTCACTGCCTCAGCCTTGGAGAG<br>CCGACGTGACCTACGCCGCCATGGTGGTGAAAGTGATCGCCCAGCACCAGAA<br>CCTGCTGCTGGCCAACACCACCAGCGCCTTCCCTTACGCCCTGCTGAGCAACG<br>ACAACGCCTTCCTGAGCTACCACCCCCACCCCTTCGCCCAGAGAACCCTGACC<br>GCCCGGTTCCAGGTCAACAACACCAGACCCCCCCACGTGCAGCTGCTGAGAA<br>AGCCCGTGCTGACCGCCATGGGACTGCTGGCCCTGCTGGACGAGGAACAGCT<br>GTGGGCCGAGGTGTCCCAGGCCGGCACCGTGCTGGACTCCAATCACACAGTG<br>GGCGTGCTGGCTAGCGCCCACAGACCTCAGGGACCCGCCGATGCTTGGCGGG<br>CTGCCGTGCTGATCTACGCCAGCGACGACACCAGAGCCCACCCCAACAGATC<br>CGTGGCCGTGACCCTGCGGCTGAGAGGCGTGCCACCTGGCCCTGGCCTGGTG<br>TACGTGACCAGATACCTGGACAACGGCCTGTGCAGCCCCGACGGCGAATGGC<br>GCAGACTGGGCAGACCTGTGTTCCCCACCGCCGAGCAGTTCCGGCGGATGAG<br>AGCCGCTGAGGACCCTGTGGCTGCCGCCCCTAGACCTCTGCCTGCTGGCGGCA<br>GACTGACCCTGAGGCCCGCTCTGAGACTGCCTTCTCTGCTGCTGGTGCACGTG<br>TGCGCCAGGCCCGAGAAGCCTCCCGGCCAGGTCACAAGACTGAGAGCCCTGC<br>CTCTGACCCAGGGACAGCTGGTGCTGGTCTGGTCCGATGAGCACGTGGGCAG<br>CAAGTGCCTGTGGACCTACGAGATCCAGTTCAGCCAGGACGGCAAGGCCTAC<br>ACCCCCGTGTCCCGGAAGCCCAGCACCTTCAACCTGTTCGTGTTCAGCCCCGA<br>CACTGGCGCTGTGTCCGGCTCTTATAGAGTGCGGGCCCTGGACTACTGGGCCA<br>GACCCGGCCCTTTCAGCGACCCCGTGCCCTACCTGGAAGTGCCCGTGCCTAGA<br>GGCCCCCCTAGCCCCGGCAACCCTCTGCGCAAGCTCCGTAAGCGGCTGCTCCT<br>GCGCAAGCTCCGTAAGCGGCTGCTCTAA |
| **eGFP** | ATGGTGAGCAAGGGCGAGGAGCTGTTCACCGGGGTGGTGCCCATCCTGGTCG<br>AGCTGGACGGCGACGTAAACGGCCACAAGTTCAGCGTGTCCGGCGAGGGCGA<br>GGGCGATGCCACCTACGGCAAGCTGACCCTGAAGTTCATCTGCACCACCGGC<br>AAGCTGCCCGTGCCCTGGCCCACCCTCGTGACCACCCTGACCTACGGCGTGCA<br>GTGCTTCAGCCGCTACCCCGACCACATGAAGCAGCACGACTTCTTCAAGTCCG<br>CCATGCCCGAAGGCTACGTCCAGGAGCGCACCATCTTCTTCAAGGACGACGG<br>CAACTACAAGACCCGCGCCGAGGTGAAGTTCGAGGGCGACACCCTGGTGAAC<br>CGCATCGAGCTGAAGGGCATCGACTTCAAGGAGGACGGCAACATCCTGGGGC<br>ACAAGCTGGAGTACAACTACAACAGCCACAACGTCTATATCATGGCCGACAA<br>GCAGAAGAACGGCATCAAGGTGAACTTCAAGATCCGCCACAACATCGAGGAC<br>GGCAGCGTGCAGCTCGCCGACCACTACCAGCAGAACACCCCCATCGGCGACG<br>GCCCCGTGCTGCTGCCCGACAACCACTACCTGAGCACCCAGTCCGCCCTGAGC |

| | |
|---|---|
| | AAAGACCCCAACGAGAAGCGCGATCACATGGTCCTGCTGGAGTTCGTGACCG<br>CCGCCGGGATCACTCTCGGCATGGACGAGCTGTACAAGTAA |
| **bGH poly(A) Signal** | CTGTGCCTTCTAGTTGCCAGCCATCTGTTGTTTGCCCCTCCCCCGTGCCTTCCT<br>TGACCCTGGAAGGTGCCACTCCCACTGTCCTTTCCTAATAAAATGAGGAAATT<br>GCATCGCATTGTCTGAGTAGGTGTCATTCTATTCTGGGGGGTGGGGTGGGGCA<br>GGACAGCAAGGGGGAGGATTGGGAAGACAATAGCAGGCATGCTGGGGATGC<br>GGTGGGCTCTATGG |
| **AAVS1 left homology arm (inverted)** | CCACTGTGGGGTGGAGGGGACAGATAAAAGTACCCAGAACCAGAGCCACATT<br>AACCGGCCCTGGGAATATAAGGTGGTCCCAGCTCGGGGACACAGGATCCCTG<br>GAGGCAGCAAACATGCTGTCCTGAAGTGGACATAGGGGCCCGGGTTGGAGGA<br>AGAAGACTAGCTGAGCTCTCGGACCCCTGGAAGATGCCATGACAGGGGGCTG<br>GAAGAGCTAGCACAGACTAGAGAGGTAAGGGGGGTAGGGGAGCTGCCCAAA<br>TGAAAGGAGTGAGAGGTGACCCGAATCCACAGGAGAACGGGGTGTCCAGGC<br>AAAGAAAGCAAGAGGATGGAGAGGTGGCTAAAGCCAGGGAGACGGGGTACT<br>TTGGGGTTGTCCAGAAAAACGGTGA |
| ***CD11b* left homology arm** | AACTTTTGGGTCTGTCATAAATAGAGGGCCCAGAATATGTAGGAGTCAGTCT<br>GGGGAGAGGCAAAGGGGATTTGGGGAAGGAGAAAGGGTTCAAGAAGAAGCA<br>GGGAGAACAGCTAGACCCAGACAGGCTGGCCAGGGAAGCCTGGATGAATGA<br>CCACATTCATGGACTGTGCAAGGCTGCTTGCCGGTCCCCTTGCTTCACACATG<br>AGGAGACGGAGGCCCAGGGAGGAGAAGTGACATGGCTCAGGGTGCGCAGCA<br>GGTGTGAGACCCCTTTCCTGAGTGCTTCCTCCTGGATCCCCTCTCACCATCTCC<br>ACTTTGCCTCCGGTTCTATTTTCCAAGGTCCCGGGTGCAAATGTTTGTTGAATG<br>ACTGATGAATGAAAATGATTTGAGTTTGTTACCTTTTATGCTTATATGTTGTGG<br>AAAATGAAATTCTCCTCAAAAGGGAAGGAAATACTTGAGAGCTGCATAGGAA<br>GGAAATTATCTAATTAAGAATGTATAGAAACTTCACTGTTGGGCAAATCATCG<br>TTGTGACACCGGGGGAAGAAGCCATTTAGGTGCTCAGAAGGGAGGCTGGAAT<br>TCAGAGCAGGACTGGACGTGCCCCACGACGGTGGTTCTTAGGTCAGGAGTCA<br>GCAAACAGTGGCCTGGGGGCCCGATATGGCCCACGACCTGTTTTTGCACAAC<br>CTGCCAGCTAGAGATTGAAGATGAACACTGATAATCGATTTGATGATAGGGA<br>GCACCACCCCCAAAGAATTCTATTTGTCTCATTTGTAAACCCGTATTACAAAC<br>AAATTGTACTCAATCATTATGTTTGAAATTTCCCTAATGACAAATTTGTGGAA<br>AAGTATTTTCTGTCTTGTTATATAAGTACTTGTACAACATATTCTATCAGCCTC<br>TT |
| **3' splicing sequence** | ATTGGCGATTTTCTTTTTAGGGC |
| **GSG linker (+2 for code frame recovery)** | CCGGAAGCGGA |

| | |
|---|---|
| **P2A** | GCTACTAACTTCAGCCTGCTGAAGCAGGCTGGAGACGTGGAGGAGAACCCTGGACCT |
| **T2A** | GAGGGCAGAGGCAGCCTGCTGACCTGCGGCGACGTCGAGGAGAACCCCGGGCCC |
| **Re-encoded *CD11b* first exon** | ATGGCTCTCAGAGTCCTTCTGTTAACA |
| **5' splicing sequence** | GGTAGGT |
| ***CD11b* right homology arm** | GGTCTGCAAAACCTAAAATTTACTATCTGGCTGTTTACAGAATAAGTGTGCTAATCCCCGCCCCAGGCTAACAGAGCTGGACCTGGGAGGCAGACATCTGGATGCTGGGTTAGTTAGGGTGACCGAATGGATGGGAAAGGGAATGGAGCAGGAAGACATGCTGCTATCTTTTTTTTTTTTTTTTTTTTTTGATACAGGGTCTTTCTCTGTTGCCCAGGCTGTAGTGCAGTGGCATGATCATGGTTCACTGCAGCCTTGACCTCCTGGGTTCAAGCAATCCTCCCACCTCAGCCTCCTGAGTACCACTACACCCGGCTAATTTTTTATTTTTTGTAGAGATGGGGTCTCACTGTGTTGCCTAGGCTGGTCTTAAACTCCTGAGCCCAGGTGATCCTCCCACGTCAGCCTCTTAAATTATTGGGATAACAGG |
